# Hierarchical assembly of the MLL1 core complex within a biomolecular condensate regulates H3K4 methylation

**DOI:** 10.1101/870667

**Authors:** Kevin E.W. Namitz, Song Tan, Michael S. Cosgrove

## Abstract

The enzymes that regulate histone H3 lysine 4 (H3K4) methylation are required for cellular differentiation and development and are often mutated in human disease. Mixed Lineage Leukemia protein-1 (MLL1) is a member of the SET1 family of histone H3 lysine 4 methyltransferases, which require interaction with a conserved sub-complex consisting of <u>W</u>DR5, <u>R</u>bBP5, <u>A</u>sh2L and <u>D</u>PY30 (WRAD_2_) for maximal activity. It is currently unclear how assembly of SET1 family complexes is involved in the spatiotemporal control of H3K4 methylation in eukaryotic genomes. In this investigation, we systematically characterized the hydrodynamic and kinetic properties of a reconstituted human MLL1 core complex and found that its assembly is highly concentration and temperature dependent. Consistent with a hierarchical assembly pathway, we found that the holo-complex assembles through interactions between the MW and RAD_2_ sub-complexes, which is correlated with enzymatic activity. Surprisingly, we found that the disassembled state is favored at physiological temperatures, and that this thermodynamic barrier can be overcome under conditions that induce high-local concentrations of subunits in phase separated compartments. Combining this data with the observation that MLL1 primary sequence contains large regions of intrinsic disorder, we propose a “swinging-domain” model in which the interaction between a tethered MW subcomplex and multiple nucleosome-RAD_2_ complexes is regulated by the rapid formation or dissolution of biomolecular condensates, such as occurs in transcription factories. This model provides an elegant “switch-like” mechanism for spatiotemporal control of H3K4 methylation within eukaryotic genomes.

## INTRODUCTION

Cellular identity in multicellular organisms is maintained in part by enzymes that regulate the degree of histone H3 lysine 4 (H3K4) methylation (1). Di- and trimethylation of H3K4 (H3K4me2,3) are enriched in gene bodies and promoters of active genes (2–4) respectively, and function to recruit nucleosome-remodeling complexes that regulate transcription (5–9). H3K4 monomethylation (H3K4me1) is associated with active gene enhancers (10–12), but is also associated with gene silencing (13–17). Because genome-wide alterations in the patterns of H3K4 methylation are linked to the aberrant transcriptional programs in developmental disorders and cancers (18–28), there is significant interest in understanding how different H3K4 methylation states are established and maintained.

Mixed Lineage Leukemia protein-1 (MLL1, ALL1, HRX, KMT2C) is a member of the SET1 family of H3K4 methyltransferases and is frequently altered in poor prognosis acute leukemias (29). MLL1 is a large protein with 3,969 amino acids and assembles into a super-complex with ~30 subunits (30–33). Subunits shared among all SET1 family members include <u>W</u>DR5, <u>R</u>bBP5, <u>A</u>sh2L, and two copies of <u>D</u>PY-30 (WRAD_2_), which associate into a sub-complex that interacts with the C-terminal SuVar, Ez, Trx (SET) domain of MLL1 (34–38). *In vitro* studies have shown that the MLL1 SET domain predominantly catalyzes H3K4 monomethylation (36), whereas multiple methylation depends on interaction of MLL1 with WRAD_2_, forming what is known as the MLL1 core complex (also known as human COMPASS, or MWRAD_2_) (34,36,39). The requirement of full MWRAD_2_ complex for optimal enzymatic activity suggests that H3K4 methylation may be regulated at the level of subunit assembly in the cell. Consistent with this hypothesis, genome-wide studies show that, while MLL1 localizes to thousands of genes in mammalian genomes, multiple methylation of H3K4 is mainly correlated with the subset of genes where MLL1 co-localizes with WRAD_2_ subunits (40). In addition, disease-specific missense mutations have been shown to disrupt MLL family core complexes (41), suggesting that aberrations in complex assembly may be associated with human disease. More recently, several labs have shown that perturbation of MLL1 core complex assembly with protein-protein interaction inhibitors may have utility as a novel therapeutic approach for treating malignancies (42–44). Together, these results suggest that knowledge of the molecular mechanisms controlling MLL1 core complex assembly will be crucial for understanding of how different H3K4 methylation states are regulated in mammalian genomes. However, progress has been impeded by the lack of understanding of the biophysical and thermodynamic mechanisms that underlie MLL1 core complex assembly.

Biochemical reconstitution studies using a minimal MLL1 SET domain construct show that the stoichiometry of the MLL1 core complex consists of one copy of the MLL1, WDR5, RbBP5 and Ash2L subunits, and 2 copies of the DPY-30 subunit (MWRAD_2_)-forming a complex with a mass of ~205 kDa (36). Direct interactions have been observed between MLL1 and WDR5 (35,37,45), WDR5 and RbBP5 (46,47), RbBP5 and Ash2L (36), and Ash2L and DPY30 (36,48,49). While these pairwise interactions suggest a linear arrangement of subunits, several lines of evidence indicate a more intricate quaternary structure. For example, while MLL1 does not interact with RbBP5 or Ash2L in pairwise experiments (36), an investigation of SET domain-associated Kabuki syndrome missense mutations suggests a direct interaction with the RbBP5/Ash2L heterodimer within the context of the holo-complex (41). The WDR5 subunit functions to stabilize this interaction by directly binding to the MLL1 <u>W</u>DR5 <u>in</u>teraction (Win) motif (35,37,45) and RbBP5 (34,36). Binding experiments show that the weakest pairwise interaction occurs between the WDR5 and RbBP5 subunits (36), suggesting the complex may be hierarchically assembled. All of these interactions have been confirmed in recent Cryo-EM and X-ray crystal structures of related SET1 family complexes (50–53). Together, these results suggest that complex assembly is hierarchical in nature, with the requirement for the formation of distinct sub-complexes before assembly of the higher-order quaternary structure. The choreographic details of this assembly pathway are unknown.

In this investigation, we systematically characterized the hydrodynamic and kinetic properties of a reconstituted human MLL1 core complex under a variety of conditions. We found that MLL1 core complex assembly is highly concentration and temperature dependent. Consistent with the hypothesized hierarchical assembly pathway, we found that the holo-complex assembles through interactions between the MW and RAD_2_ sub-complexes, and that MWRAD_2_ formation is directly correlated with enzymatic activity. Surprisingly, we found that the disassembled state is favored at physiological temperatures and at concentrations typically used in steady-state enzymatic assays. In contrast, sub-physiological ionic strength dramatically increases enzymatic activity, which is associated with the formation of induced high-local concentrations of the MLL1 core complex in phase-separated droplets. Based on these results, we propose a model in which the thermodynamic barrier to complex assembly is overcome in the cell under conditions that induce high-local concentrations of subunits, such as those found in transcription factories. Together, these results are consistent with the hypothesis that regulated assembly of the MLL1 core complex underlies an important mechanism for establishing different H3K4 methylation states in mammalian genomes.

## RESULTS

### MLL1 core complex assembly is concentration and temperature dependent

To better understand MLL1 core complex assembly, we purified human recombinant MWRAD_2_ as described in Methods and characterized its oligomeric behavior by size exclusion chromatography (SEC) and sedimentation velocity analytical ultracentrifugation (SV-AUC). SEC revealed that the purified complex eluted as a single symmetrical peak (Fig. 1A) and SDS-PAGE of the indicated fractions showed the presence of all five subunits with the expected stoichiometry (Fig. 1B). We note that the complex elutes later than expected based on its theoretical mass, which is likely due to the significant shape asymmetry of the particle. We then chose SV-AUC to characterize the concentration and temperature dependence of the complex in solution. SV-AUC is a first-principle technique that measures the time course of sedimentation of macromolecules in a gravitational field in a way that maintains the equilibrium of reversible associations – allowing extraction of equilibrium and kinetic properties of interactions (54,55). Sedimentation boundaries formed as the particle sediments over time were fit using a finite element analysis of Lamm equation solutions (Fig. 1C) (56) to give the diffusion-free sedimentation coefficient distribution *c(s)* (Fig. 1D). The *c(s)* plot of MWRAD_2_ at 5 μM loading concentration at 5°C revealed a large peak accounting for almost 90% of the signal with an *s*_*20,w*_ (*S*) value of 7.2 and two minor peaks at 2.9 and 4.7 *S* that each account for 4-5% of the signal (noted with arrows in Fig. 1D). The major peak at 7.2 *S* corresponds to the fully assembled MLL1 core complex, which we previously showed assembles with a stoichiometry of 1:1:1:1:2 for the MWRAD_2_ subunits, respectively (36). In addition, the *S*-value of MWRAD_2_ is independent of loading concentration (Fig. 2A), indicating that the complex is stable at 5°C and has a relatively long lifetime compared to the timescale of sedimentation (57). Using the derived weight-averaged frictional coefficient (*f/f*_*0*_) of 1.7, the calculated molecular mass from this *S* value was 209,561 Daltons, which is within error of the expected mass (205,402) based on the amino acid sequence of the holo-complex subunits at the indicated stoichiometry.

**Figure 1:**
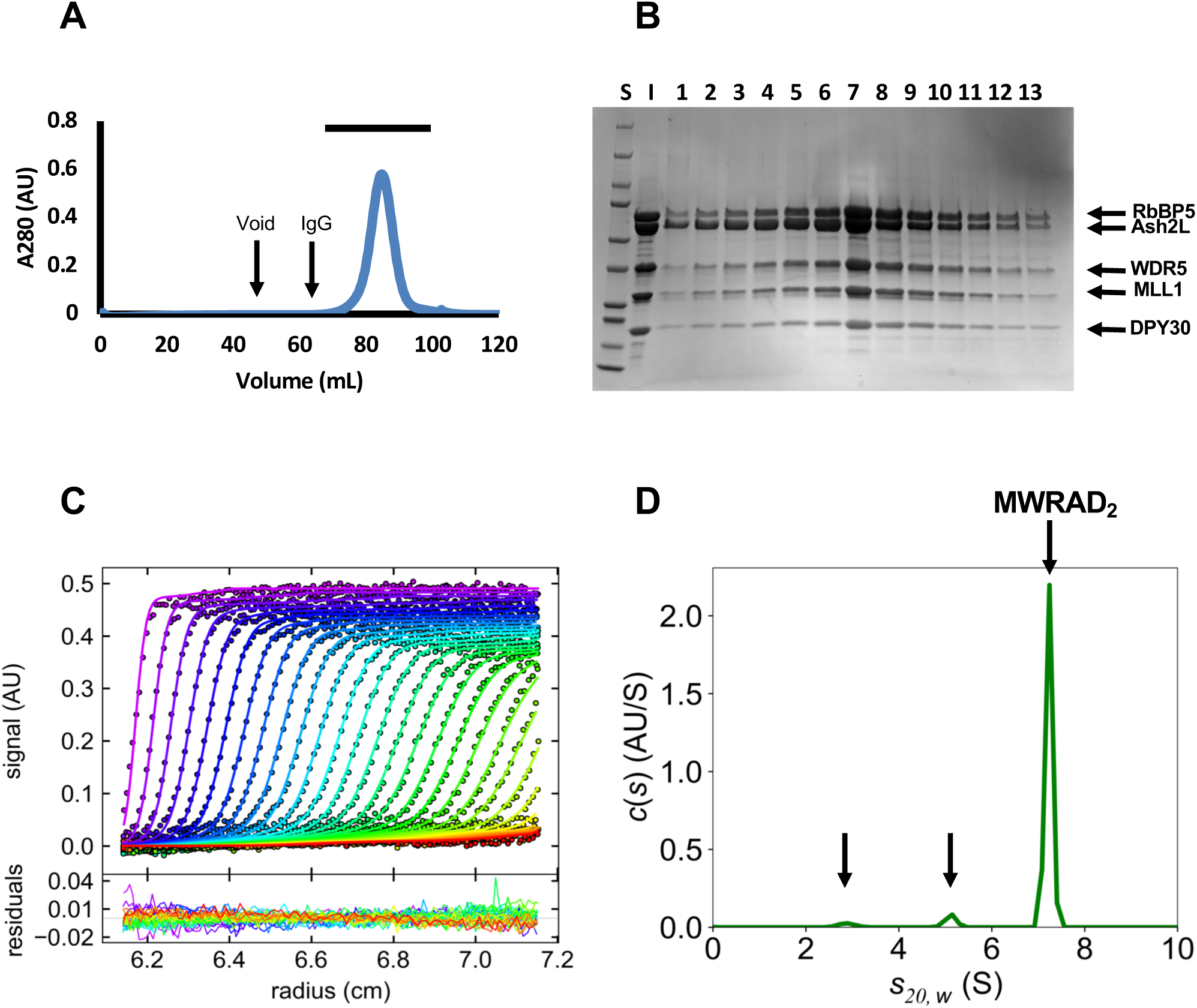
Purification and characterization of the MLL1 core complex. (A) Chromatogram of S200 SEC purified MWRAD_2_. The void volume and elution volume of IgG (M_r_ 158 kDa) are indicated. The horizontal bar above the peak indicates fractions shown on the Coomassie stained SDS-PAGE gel in (B). (C) Upper Panel: SV-AUC run showing raw data (points) and fits using the continuous distribution (*c(s)*) method by the program SEDFIT (solid lines) (56). The lower panel shows the residuals derived from the fit. Shown is a typical run of 5 μM MWRAD_2_ taken at 5°C. (D) Diffusion-deconvolved sedimentation coefficient distribution (*c(s)*) obtained using the fits to the raw data shown in (C). All profiles are shown with experimental *s\** values corrected to standard conditions at 20°C in water (*s*_*20,w*_ (*S*)). The positions of MWRAD_2_ and the two minor peaks are indicated with arrows.

**Figure 2:**
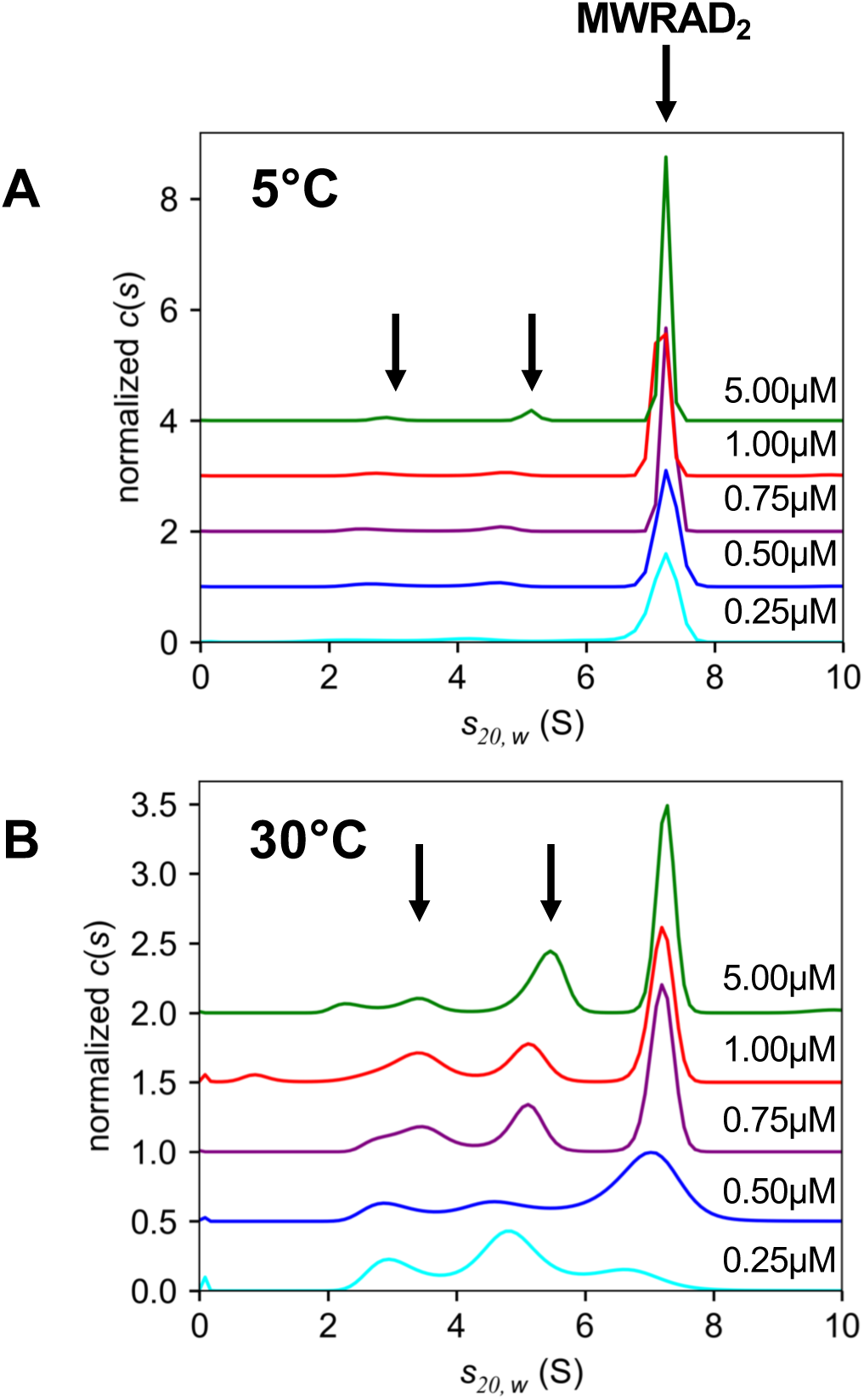
The holo-MLL1 core complex assembles from predominantly two sub-complexes. (A) *c(s)* distributions of MWRAD_2_ at 5°C at five different concentrations: 0.25 μM (cyan), 0.5 μM (blue), 0.75 μM (purple), 1.0 μM (red) and 5.0 μM (green). Each profile was normalized by total integrated area under the peaks. (B) The same as in (A), but at 30°C. The unlabeled arrows in both (A) and (B) indicate the positions of the putative sub-complex peaks at the highest concentration.

The minor peaks observed in the *c(s)* distribution in Fig. 1D could represent trace contaminants in the sample or minor populations of dissociated sub-complexes and/or subunits. To distinguish these hypotheses, we compared *c(s)* distributions of MWRAD_2_ at concentrations ranging from 0.25 – 5 μM at 5°C (Fig. 2A) and 30°C (Fig. 2B). If the minor peaks represent non-interacting contaminants, then the relative amount of signal between the major and minor peaks will not vary as the loading concentration is decreased. In contrast, if the complex is dissociating into sub-complexes, then the relative amount of signal in the major and minor peaks will change as the loading concentration is varied. The results were consistent with the latter possibility. For example, while the effect at 5°C was modest, when the loading concentration of the complex was decreased from 5 μM to 0.25 μM, the amount of signal corresponding to the holo-complex decreased from ~88% to ~83% of the total signal, with a corresponding increase in both minor peak signals (Fig. 2A). The effect was more obvious at 30°C, which showed that the signal corresponding to the minor peaks increased from 35% to 75% of the total signal as the loading concentration was decreased (Fig. 2B). These results suggest that the minor peaks represent dissociated sub-complexes and/or subunits. Furthermore, because the *S*-values of the minor peaks show varying degrees of concentration dependence, they likely represent reaction boundaries of sub-complexes as opposed to individual non-interacting subunits. These data suggest that the holo-complex assembles from predominantly two sub-complexes in a temperature and concentration-dependent manner.

### The disassembled state of the MLL1 core complex is favored at physiological temperature

To further explore the thermodynamics of MLL1 core complex assembly, we compared the temperature dependence of MWRAD_2_ formation at several different loading concentrations using SV-AUC (Fig. 3). Each *c(s)* profile was integrated and the relative amount of signal corresponding to the *S* value of the holo-complex was plotted as a function of temperature and total loading concentration (Fig. 3F). At the highest loading concentration (5 μM), little variation in the amount of holo-complex was observed between 5° and 25°C (Fig. 3A, F), with a peak that accounted for 81-92% of the total signal (Table S1). In contrast, at temperatures greater than 25°C, the amount of holo-complex decreased precipitously until only ~3% of the signal could be observed at 37°C (Figs. 3A and F, Table S1). The effect of temperature on MLL1 core complex stability became increasingly more severe as the loading concentration was decreased. For example, at the lowest loading concentration (0.25 μM), only the 5°C and 10°C runs showed ~80% holo-complex (Fig. 3E, F; Table S1); whereas at higher temperatures, the signal corresponding to the holo-complex decreased from ~63% at 15°C - to ~2% of the total signal at 37°C (Fig. 3F; Table S1). At 37°C, most of the signal is instead dominated by the two sub-complex peaks with *S*-values of ~3 and 4.7 (Fig. 3G). These data are consistent with the hypothesis that the holo-MLL1 core complex assembles from interaction of two sub-complexes, the equilibrium of which is highly concentration and temperature-dependent.

**Figure 3:**
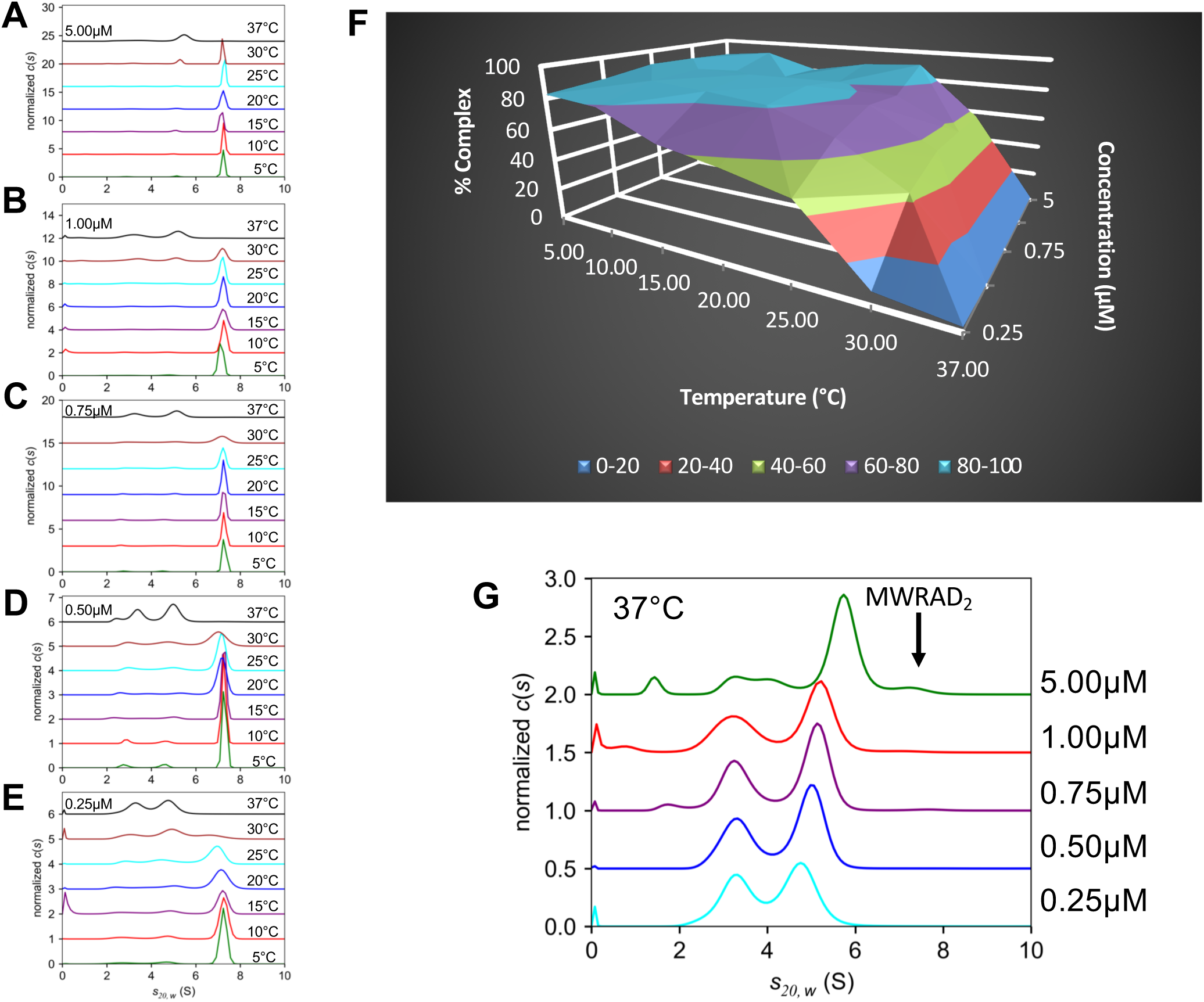
Temperature dependence of MLL1 core complex assembly. (A-E) Representative *c(s)* distributions of the MLL1 core complex at the indicated temperatures and loading concentrations. Each distribution was normalized for total integrated area. (F) Surface plot summarizing the percentage of signal in SV-AUC experiments corresponding to the *S* value of the MLL1 core complex as a function of temperature and concentration (see also Table S1). These values were obtained as described in Methods. (G) *c(s)* distributions from five MWRAD_2_ concentrations at 37°C normalized by total integrated area (note: each distribution corresponds to the black line from the respective concentration panel in A – E). The position of holo-MWRAD_2_ at 7.2 *S* is indicated with the arrow.

Surprisingly, at all the concentrations tested, very little holo-complex with an *S* value of 7.2 was observed at physiological temperature (37°C) (Fig. 3G). This suggests that the disassembled state of the MLL1 core complex may predominate in cells, and that other factors are required to stabilize the assembled state. In support of this hypothesis, closer examination of the *c(s)* profiles of the complex at 37°C revealed evidence that increased protein concentration promotes complex formation. For example, while similar amounts of signal are observed in the two sub-complex peaks at the 0.25 μM loading concentration (cyan line, Fig. 3G), the relative amount of signal in the two peaks changes with progressively higher concentrations. The intensity of the larger peak increased at the expense of the smaller peak and began to show evidence of concentration-dependent shifting to higher *S*-values. This hydrodynamic behavior is consistent with a reaction boundary composed of free and bound reactants that interconvert under a rapid kinetic regime that cannot be resolved within the signal-to-noise of the experiment (57). These results suggest that, unlike the long lifetime of the assembled complex observed at 5°C, the kinetics of the interaction have changed at 37°C such that the complex now has a short lifetime compared to the timescale of sedimentation.

We next analyzed the concentration series at each temperature to derive binding isotherms. We integrated each *c(s)* profile (between 0.5 and 9.5 *S*) to determine the weight-average sedimentation coefficients (*s_w_*) (58), which were then plotted against MWRAD_2_ concentration and fit to derive the apparent dissociation constant 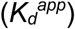 for each isotherm (Fig. 4A). Given that the majority of signal in each *c(s)* profile could be assigned to three peaks, we applied the A + B ⇆ AB hetero-association model in the program SEDPHAT (59) and obtained reasonable fits (Table 1). The derived 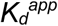 values ranged from 7 nM at 5°C to ~6,200 nM at 37°C (Table 1). A van’t Hoff analysis showed that complex formation is exothermic, which is offset by the negative entropy change as the complex subunits become more ordered (Fig. 4B and C). However, the van’t Hoff plot reveals a non-linear relationship between K_eq_ and temperature, indicating a change in the heat capacity of the system at higher temperatures (Fig. 4B). These data suggest at least two mechanisms for complex assembly, which differ by temperature. At low temperatures (≤ 25°C), the equilibrium favors complex formation, with a relatively long lifetime that is stable on the timescale of sedimentation. Under this mechanism, the interaction is dominated by enthalpic contributions to the free energy (Fig. 4C). At high temperatures (> 25°C), the equilibrium is shifted into the rapid kinetic regime with a short complex lifetime where dissociation is more likely. While there is little difference in the Gibbs free energy between mechanisms, there is a difference in the contributions between the enthalpic and entropic terms. At higher temperatures, the entropic penalty to complex formation was increased 7-fold compared to that of the lower temperature mechanism, while the difference in the enthalpic contribution was only increased by 3.8-fold (Fig. 4C). These results suggest that, at physiological temperature, one or more of the subunits samples alternate conformational states, some of which are not competent for complex assembly. However, given the observation that some holo-complex forms in a concentration-dependent manner, increased local concentration of subunits may be a mechanism that cells use to overcome the increased entropic cost of complex formation at 37°C.

**Table 1:** Summary of apparent dissociation constants for MLL1 core complex assembly at different temperatures*

| Temperature (°C) | $K_d^{app}$ (nM) | Confidence interval (1σ) |
| --- | --- | --- |
| 5 | 7 | 6 – 9 |
| 10 | 7 | 5 – 11 |
| 15 | 20 | 13 – 30 |
| 20 | 30 | 22 – 39 |
| 25 | 62 | 51 – 72 |
| 30 | 290 | 204 – 417 |
| 37 | 6200 | 4900 – 7900 |

\* Dissociation constants and error estimates were obtained from fitting MWRAD<sub>2</sub> concentration versus signal weight average sedimentation coefficient ( $s_w$ ) using the $A + B \rightleftharpoons AB$ hetero-association model in SEDPHAT (59).

**Figure 4:**
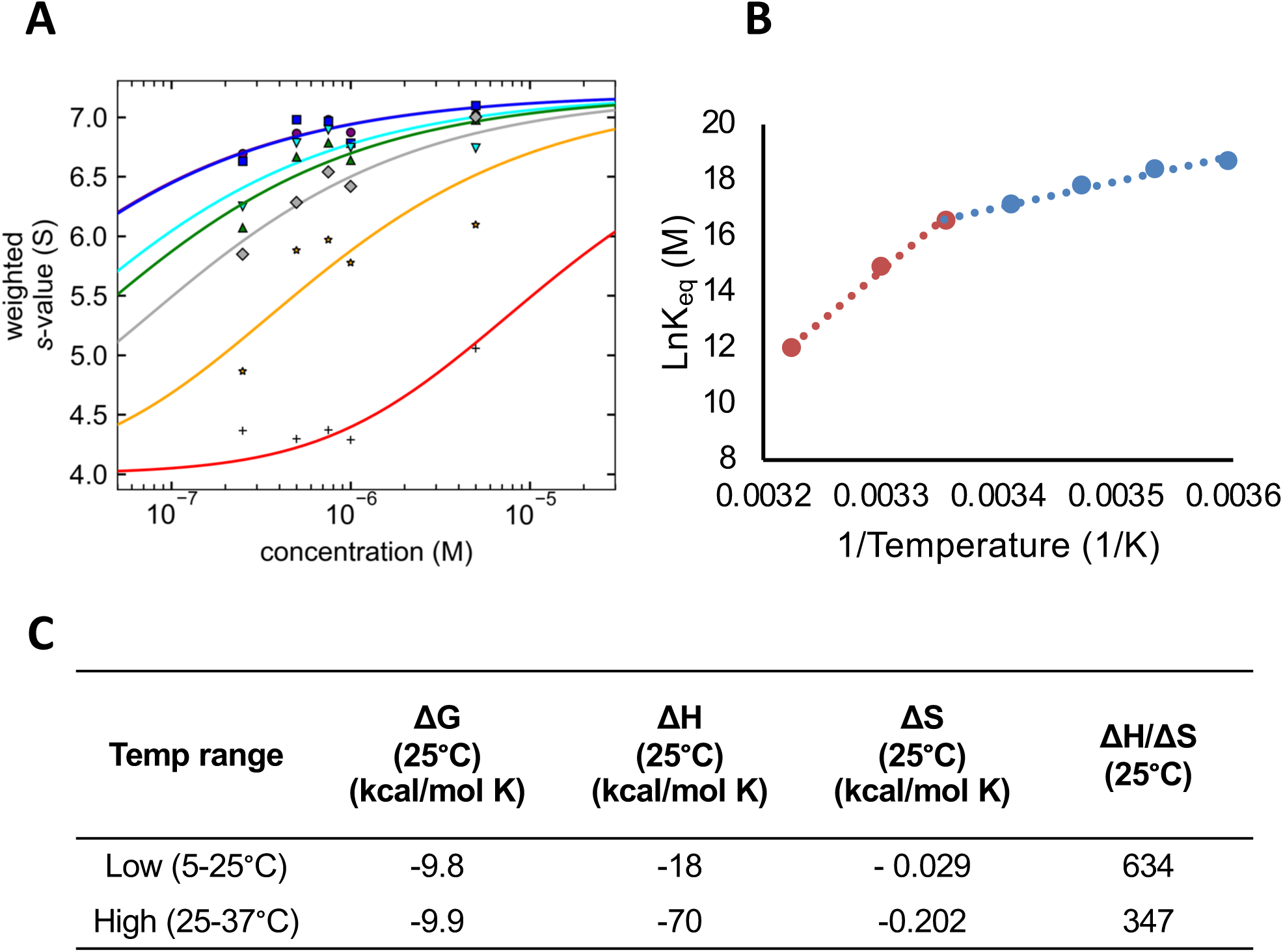
Thermodynamic characterization of MLL1 core complex assembly. (A) Signal-weighted (*s*_*w*_) isotherms of MWRAD_2_ were obtained for each temperature, plotted against loading concentrations and fit to an A + B ⇋ AB hetero-association model using SEDPHAT (114). The lines represent the fits for each isotherm, which were conducted at 5°C (blue),10°C (purple), 15°C (cyan), 20°C (green), 25°C (grey), 30°C (orange) and 37°C (red). 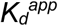 values are summarized in Table 1. (B) van’t Hoff plot derived from the apparent *K*_*eq*_ values. Linear regression was used to independently fit the data for the high temperature range (red, 25-37°C) and low temperature range (blue, 5-25°C). (C) Summary of thermodynamic parameters for MLL1 core complex assembly under high and low temperature regimes derived from the van’t Hoff analysis in (B).

### The MLL1 core complex assembles from MW and RAD_2_ sub-complexes

Previous experiments suggested that the holo-complex is assembled by pairwise interactions as follows: M ⇌ W ⇌ R ⇌ A ⇌ D_2_ (36). Since the weakest pairwise interaction occurs between WDR5 and RbBP5 (36), we predicted that the complex assembles by first forming MW and RAD_2_ sub-complexes, which then interact to form the holo-complex (Scheme 1). However, we reasoned that there are at least two additional reaction schemes that could potentially give rise to the three boundaries observed in the holo-complex *c(s)* profiles (Schemes 2 and 3). To distinguish among these schemes, we chose to use a Bayesian approach to analyze the SV-AUC data of the holo-complex collected at 25°C. The Bayesian approach is a variant of the standard maximum entropy regularization method utilized in the *c(s)* analysis in that, instead of assuming a uniform probability for the occurrence of species at every *S*-value in a distribution, it utilizes prior information to assign different probabilities in different regions of *S*-values (60). A key feature of the Bayesian implementation in SEDFIT is that, because it maintains the same degrees of freedom used in the standard *c(s)* analysis, imperfections in the expected values will result in additional features in the *c*^(*p*)^(*s*) plots in order to maintain the quality of the fit (60). The Bayesian analysis therefore allows us to determine which reaction scheme gives a *c*^(*p*)^(*s*) profile that best fits the experimental data.

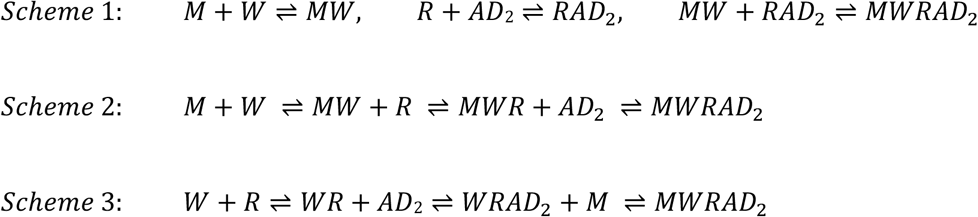

To obtain the expected *S*-values for each of the predicted sub-complexes or subunits in each reaction scheme, we mixed stoichiometric amounts of their respective subunits and characterized their concentration dependence by SV-AUC at 25°C (Fig. S1; Table S2). We then used each of the *S*-values collected at 0.25 μM as prior expectations in the Bayesian analysis of the holo-complex. As shown in Fig. 5A, when the independently determined *S*-values for MW, RAD_2_ and the MWRAD_2_ species were used as prior expectations in the Bayesian analysis of the holo-complex at 0.25 μM (black dotted line), three peaks in the *c*^(*p*)^(*s*) plot were observed that were in excellent agreement with the expectations (cyan line). Indeed, good agreement was observed using the same *S*-values as prior expectations for Bayesian fits of the experimental data collected at higher holo-complex concentrations (Fig. 5A). The only deviation observed was for the position and amplitude of the holo-complex peak, which at 25°C shifts from 6.8 to 7.2 *S* in a concentration-dependent manner (Fig. 5A). In contrast, when a similar analysis was conducted instead using the expected *S*-values for the MWR and AD_2_ sub-complexes predicted by Scheme 2, additional features in the *c*^(*p*)^(*s*) plot with an *S*-value of ~5.3 were observed at all loading concentrations that did not match the prior expectations (Fig. 5B, red arrow). Similarly, using the expected *S*-values for M and WRAD_2_ as predicted by Scheme 3, the *c*^(*p*)^(*s*) plot showed little evidence of a species matching the expected value of free MLL1 at 2.3 *S*, and also showed additional features at ~3.5 *S* that did not match expectations (Fig. 5C, red arrow). To test whether the holo-complex assembles in a concerted fashion from individual subunits, we also performed a similar Bayesian analysis using the predetermined *S* values for M, W, R, AD_2_, and MWRAD_2_ as prior expectations (AD_2_ is treated as a discrete species since it does not appreciably dissociate under the range of concentrations that can be detected by the absorbance optical system used in these experiments (36)). The *c*^(*p*)^(*s*) plot showed additional features with an *S*-value of ~5.2 that did not match expectations (Fig. 5D, red arrow). Together, these results are consistent with the hypothesis that MLL1 core complex is hierarchically assembled by association of MW and RAD_2_ sub-complexes.

**Figure 5:**
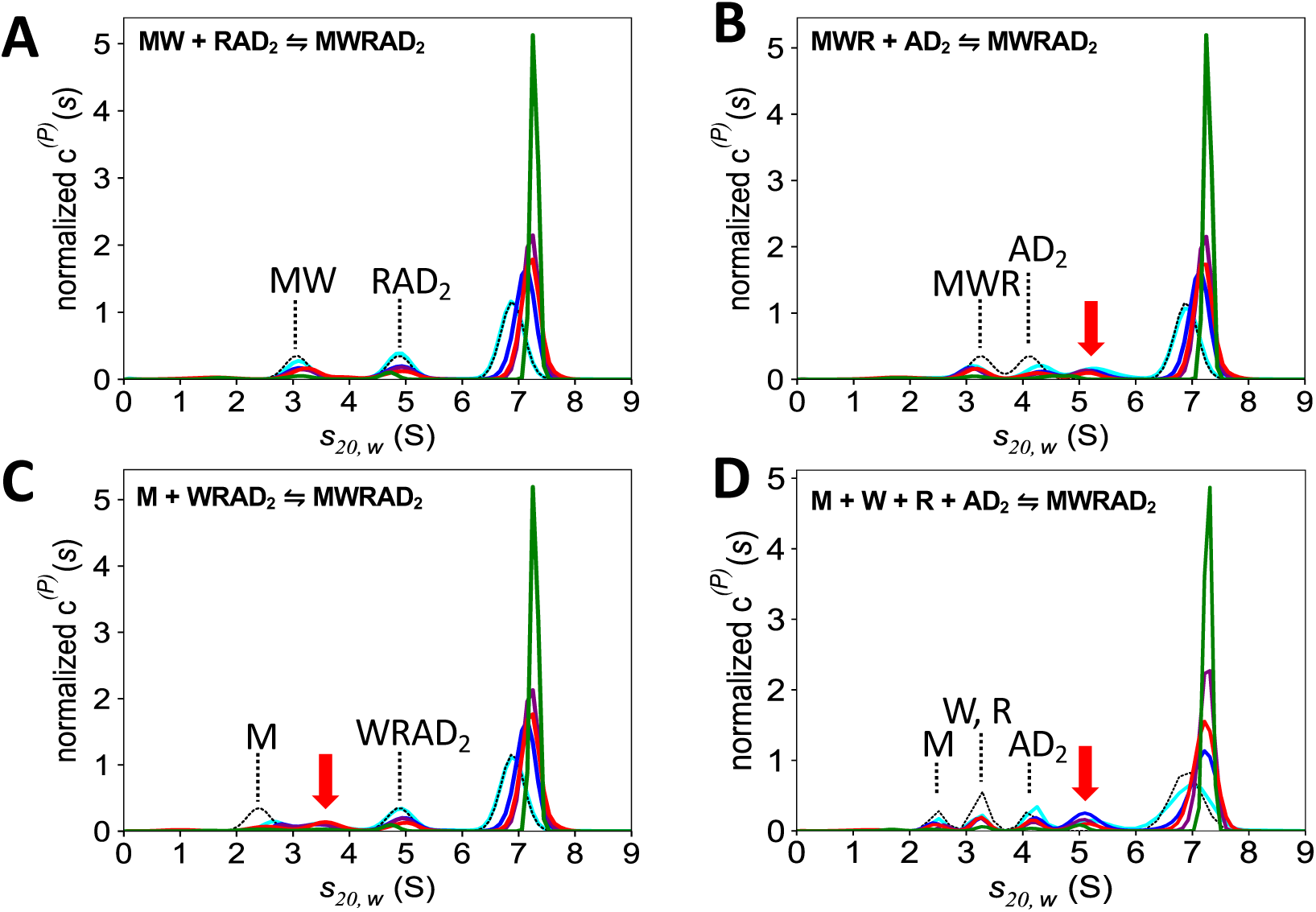
The holo-MLL1 core complex assembles from MW and RAD_2_ sub-complexes. Bayesian analysis of MWRAD_2_ SV-AUC data collected at 25°C. MWRAD_2_ concentrations were 0.25 μM (cyan), 0.5 μM (blue), 0.75 μM (purple), 1.0 μM (red) and 5.0 μM (green). Maximum entropy regularizations were restrained with expected values (indicated with the dotted line) for each indicated sub-complex derived from separate experiments (Fig. S1 and Table S2) to give *c*^(*p*)^(*s*) distributions (colored lines), which were normalized by total integrated area. Concentrations of MWRAD_2_ in each run were: 0.25 μM (cyan), 0.5 μM (blue), 0.75 μM (purple), 1.0 μM (red) and 5.0 μM (green). The *c*^(*p*)^(*s*) distributions used the following *S* values as prior expectations: (A) MW (3.0 S), RAD_2_ (4.4 S), and MWRAD_2_ (6.9 S) (scheme 1); (B) MWR (3.3 S), AD_2_ (4.1 S), and MWRAD_2_ (6.9 S) (scheme 2). (C) M (2.3 S), WRAD_2_ (4.4 S), and MWRAD_2_ (6.9 S) (scheme 3). (D) M(2.3 S), W(3.2 S), R(3.4 S), AD_2_(4.1 S), and MWRAD_2_ (6.9 S) (concerted assembly scheme).

### Enzymatic activity of the MLL1 core complex is directly related to complex assembly

To determine the impact of concentration and temperature on the enzymatic activity of the MLL1 core complex, we incubated MWRAD_2_ (0.25 – 5 μM) with a fixed concentration of histone H3 peptide (10 μM) and saturating amounts of AdoMet (250 μM) at various temperatures. We then measured methylation using a label-free quantitative MALDI-TOF mass spectrometry assay (36). MALDI spectra were integrated and the relative amount of each peptide species was plotted as a function of time. Data were fit using a numerical integration of rate equations approach implemented in KinTek Explorer software (61), which allowed us to test the ability of different reaction schemes to fit the data.

Using the simplest irreversible consecutive reactions model (Fig. 6, Scheme 4), while acceptable fits were obtained for reaction progress curves collected at the highest concentration (5 μM) between temperatures 5 – 30°C (5°C is shown in Fig. 6A), the rest of the fits were poor (an example is shown in Fig. 6B). Since we previously showed that the complex uses a non-processive mechanism for multiple lysine methylation (36), we revised the model to incorporate binding of peptide substrate to the enzyme-AdoMet complex (E_1_) and release of the H3K4me1 product after the first methylation event, followed by binding of the H3K4me1 substrate to a distinct site on the enzyme (E_2_) for the dimethylation reaction. The latter step is predicated on our previous observation that the MLL1 core complex has a cryptic second active-site independent of the SET domain that is required for the H3K4 dimethylation reaction (36,62,63). Since the binding and release rates of substrates and product are currently unknown, these values were fixed to be non-rate limiting. This model allowed us to incorporate an additional term to test the impact of reversible complex disassembly, which results in negligible activity of both enzymes under these assay conditions (Fig. 6, Scheme 5) (36,37). Initial values for the ratio (*k*_*off*_/*k*_*on*_) for complex assembly were set to be equal to the 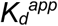 derived from each SV-AUC isotherm experiment.

**Figure 6:**
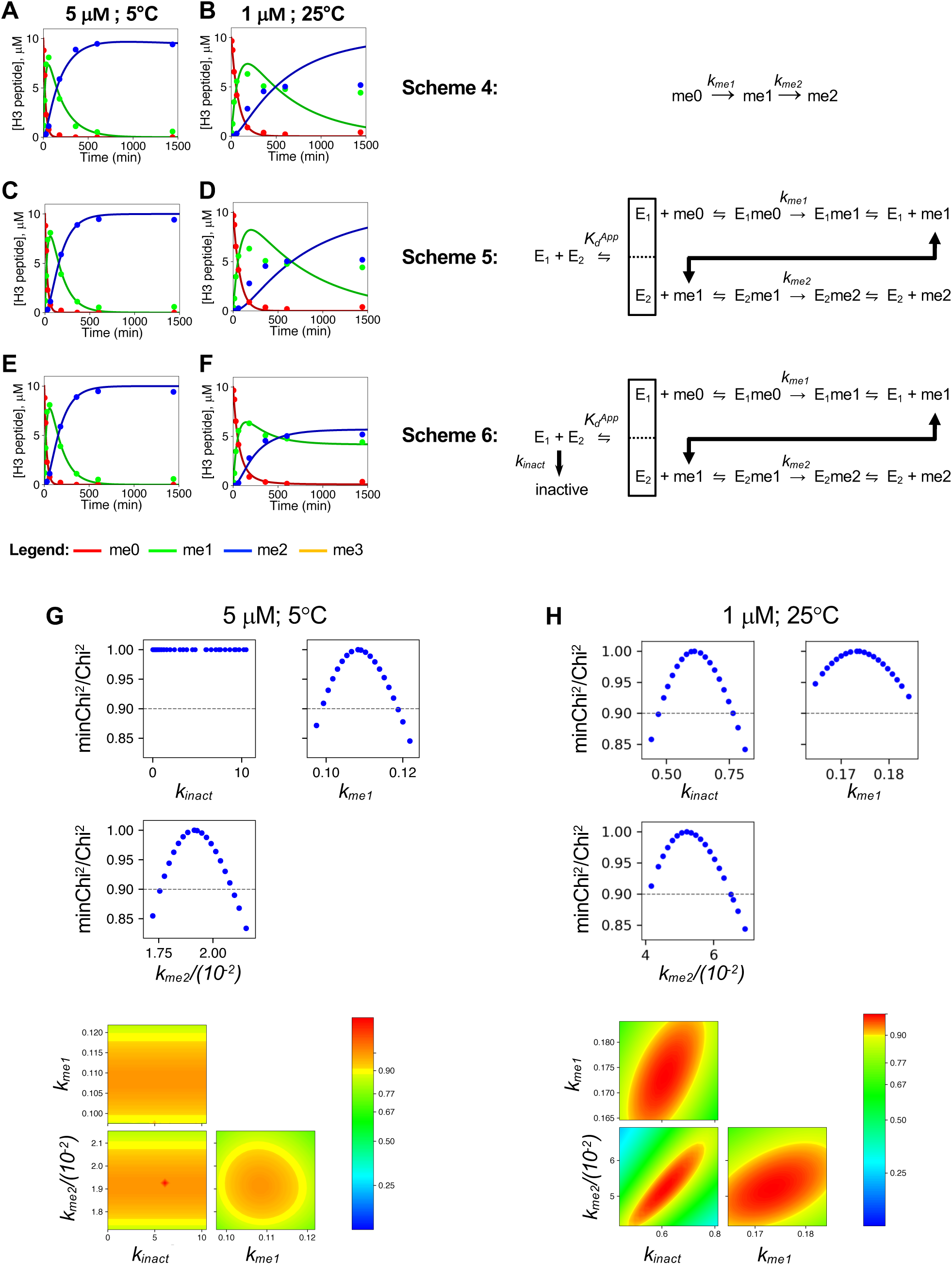
Comparison of minimal reaction pathways. (A, C, E) show the fits (solid lines) for the concentrations of each peptide species (me0, me1, or me2) during the reaction time course catalyzed by 5 μM MWRAD_2_ at 5°C. Each panel shows the fits to the same data using Scheme 4 (A), Scheme 5 (C), or Scheme 6 (E). Panels (B, D, F) show fits for Schemes 4-6, respectively, for the reaction time course catalyzed by 1 μM MWRAD_2_ at 25°C. (G) Fitspace confidence contour analysis for the reaction catalyzed by 5 μM MWRAD_2_ at 5°C fit with Scheme 6. *k*_*inact*_ is not constrained by the data, mainly due to the absence of detectable enzyme inactivation during the reaction time course at 5°C. (H) Fit space confidence contour analysis of the fit of Scheme 6 to the reaction catalyzed by 1 μM MWRAD_2_ at 25°C. *k*_*inact*_ is now constrained by the data.

The resulting simulations showed that adding a reversible complex disassembly step to the reaction scheme only modestly improved fits to the lower temperature data (Fig. 6C), but did not improve the fits of the higher temperature data (Fig. 6D). In addition, Fitspace confidence contour analysis (64) showed that the derived *k*_*off*_ value for the complex dissociation step was not constrained by the data (not shown), suggesting that the model is more complex. Closer examination of the high temperature data showed that several reactions failed to go to completion, suggesting the enzyme rapidly inactivates at higher temperatures. We therefore revised the working model to incorporate an irreversible enzyme inactivation step (*k*_*inact*_) (Figure 6, Scheme 6). The resulting simulations resulted in good fits to both the low and high temperature datasets shown in Figs. 6E and 6F, respectively. In addition, Fitspace analysis showed that the derived pseudo-first order rate constants for monomethylation (*k*_*me1*_), and dimethylation (*k*_*me2*_) reactions were reasonably well-constrained by the data (Fig. 6G and H). Furthermore, the rate of enzyme inactivation (*k*_*inact*_) was constrained by the data in the higher temperature experiments (Fig. 6H) but not in the lower temperature experiments (Fig. 6G), where enzyme inactivation is negligible. Figure 7 shows that the use of Scheme 6 produces good fits for all datasets.

**Figure 7:**
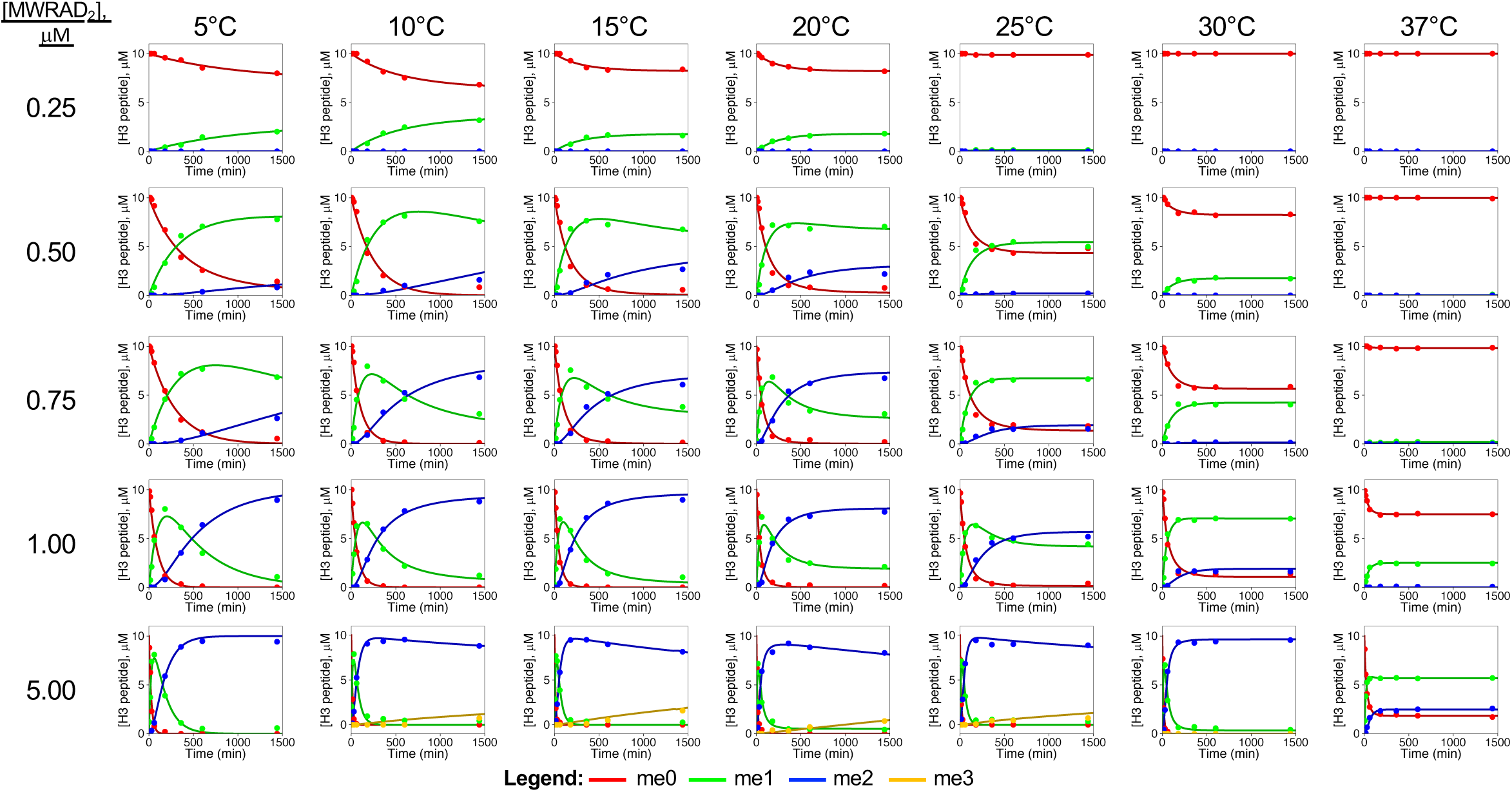
Temperature and concentration dependence of MLL1 core complex enzymatic activity. Time courses for reactions at the indicated MWRAD_2_ concentrations and temperatures were plotted and fit using Scheme 6. Each time point represents the average from two independent experiments. Concentrations of each peptide species were plotted in red for H3K4me0, green for H3K4me1, blue for H3K4me2. For reactions showing small amounts of H3K4me3 (yellow), Scheme 6 was modified to incorporate an additional turnover step followed by product release.

Based on these results, we then used the fits to Scheme 6 to compare the impacts of temperature and concentration on the enzymatic activity of the MLL1 core complex (Fig. 8). The obtained pseudo-first order rate constants for monomethylation (*k*_*me1*_), dimethylation (*k*_*me2*_) and the rate of enzyme inactivation (*k*_*inact*_) are summarized in Tables 2–4, respectively. At most of the tested enzyme concentrations, activity increased linearly as the temperature increased from 5°C to 20°C (Fig. 8A and C). However, above 20°C, non-Arrhenius behavior was observed, as the rate of irreversible enzyme inactivation (*k*_*inact*_) rivaled or exceeded the rates of turnover (Tables 2–4), resulting in reactions that failed to go to completion (Fig. 7). These results are consistent with the conclusions from the SV-AUC analysis, which suggested that as the complex dissociates at higher temperatures, one or more of the subunits undergoes an irreversible conformational change that is not competent for catalysis. We therefore plotted *k*_*me1*_ and *k*_*me2*_ rates (Ln(*k*_*n*)_) as a function of temperature (1/T) between 5°C and 20°C to fit the data to the Arrhenius equation (Fig. 8 B and D, respectively). Linear fitting of the Arrhenius plots revealed similar values for the energy of activation (*E*_*a*_) between the tested concentrations. The average *E*_*a*_ values were 10.9 ± 2.0 kcal K^−1^ mol^−1^ and 17.8 ± 4.7 kcal K^−1^ mol^−1^ for the monomethylation and dimethylation reactions, respectively.

**Table 2.** Pseudo-first order rate constants for H3K4 monomethylation (*k*_*me1*_) catalyzed by MWRAD_2_ at the indicated concentration and temperature*

| Temperature: | 5°C | 10°C | 15°C | 20°C | 25°C | 30°C | 37°C |
| --- | --- | --- | --- | --- | --- | --- | --- |
| [MWRAD<sub>2</sub>], μM | $k_{me1}, min^{-1}$ | $k_{me1}, min^{-1}$ | $k_{me1}, min^{-1}$ | $k_{me1}, min^{-1}$ | $k_{me1}, min^{-1}$ | $k_{me1}, min^{-1}$ | $k_{me1}, min^{-1}$ |
| 0.25 | 0.01 ± 0.01 | 0.03 ± 0.02 | 0.03 ± 0.04 | 0.03 ± 0.03 | 0.0 ± 0.05 | N/A<sup>a</sup> | N/A |
| 0.5 | 0.06 ± 0.01 | 0.10 ± 0.02 | 0.15 ± 0.03 | 0.18 ± 0.03 | 0.11 ± 0.05 | 0.03 ± 0.02 | N/A |
| 0.75 | 0.06 ± 0.01 | 0.16 ± 0.02 | 0.16 ± 0.02 | 0.24 ± 0.02 | 0.12 ± 0.04 | 0.07 ± 0.05 | 0.00 ± 0.02 |
| 1.0 | 0.13 ± 0.01 | 0.19 ± 0.04 | 0.25 ± 0.04 | 0.29 ± 0.04 | 0.19 ± 0.04 | 0.18 ± 0.04 | 0.07 ± N.D.<sup>b</sup> |
| 5.0 | 0.13 ± 0.02 | 0.26 ± 0.04 | 0.31 ± 0.04 | 0.32 ± 0.04 | 0.30 ± 0.04 | 0.28 ± 0.04 | 0.13 ± 0.06 |

\* Each is the rate constant +/- the Standard Error determined from duplicate measurements.
<sup>a</sup>N/A, Not applicable – no methylation observed under the indicated condition.
<sup>b</sup>N.D., error estimates are not defined.

**Table 3.**
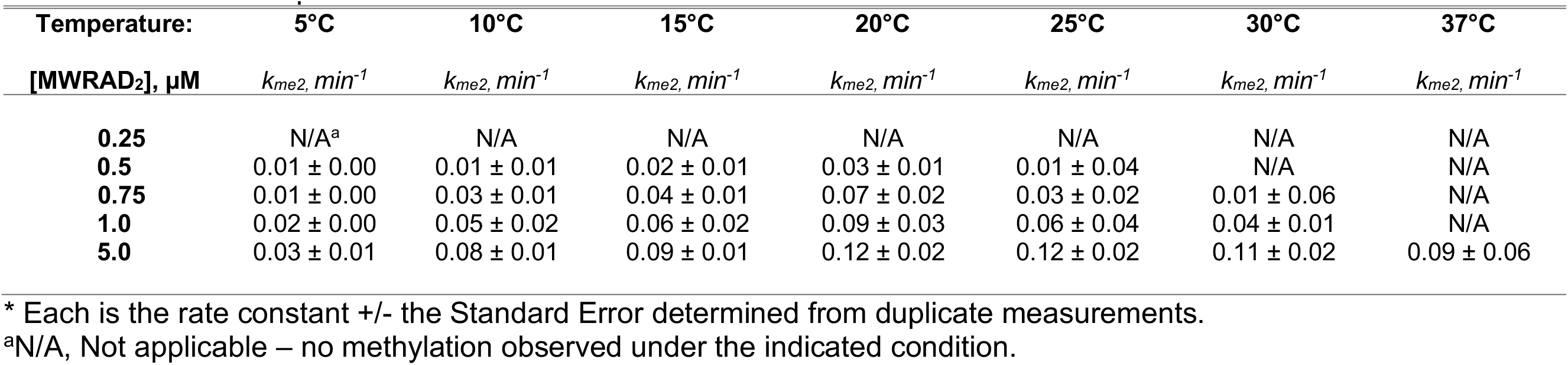
Pseudo-first order rate constants for H3K4 dimethylation (*k*_*me2*_) catalyzed by MWRAD_2_ at the indicated concentration and temperature*

| Temperature: | 5°C | 10°C | 15°C | 20°C | 25°C | 30°C | 37°C |
| --- | --- | --- | --- | --- | --- | --- | --- |
| [MWRAD<sub>2</sub>], μM | $k_{me2}, \text{min}^{-1}$ | $k_{me2}, \text{min}^{-1}$ | $k_{me2}, \text{min}^{-1}$ | $k_{me2}, \text{min}^{-1}$ | $k_{me2}, \text{min}^{-1}$ | $k_{me2}, \text{min}^{-1}$ | $k_{me2}, \text{min}^{-1}$ |
| **0.25** | N/A<sup>a</sup> | N/A | N/A | N/A | N/A | N/A | N/A |
| **0.5** | 0.01 ± 0.00 | 0.01 ± 0.01 | 0.02 ± 0.01 | 0.03 ± 0.01 | 0.01 ± 0.04 | N/A | N/A |
| **0.75** | 0.01 ± 0.00 | 0.03 ± 0.01 | 0.04 ± 0.01 | 0.07 ± 0.02 | 0.03 ± 0.02 | 0.01 ± 0.06 | N/A |
| **1.0** | 0.02 ± 0.00 | 0.05 ± 0.02 | 0.06 ± 0.02 | 0.09 ± 0.03 | 0.06 ± 0.04 | 0.04 ± 0.01 | N/A |
| **5.0** | 0.03 ± 0.01 | 0.08 ± 0.01 | 0.09 ± 0.01 | 0.12 ± 0.02 | 0.12 ± 0.02 | 0.11 ± 0.02 | 0.09 ± 0.06 |

\* Each is the rate constant +/- the Standard Error determined from duplicate measurements.
<sup>a</sup>N/A, Not applicable – no methylation observed under the indicated condition.

**Table 4.**
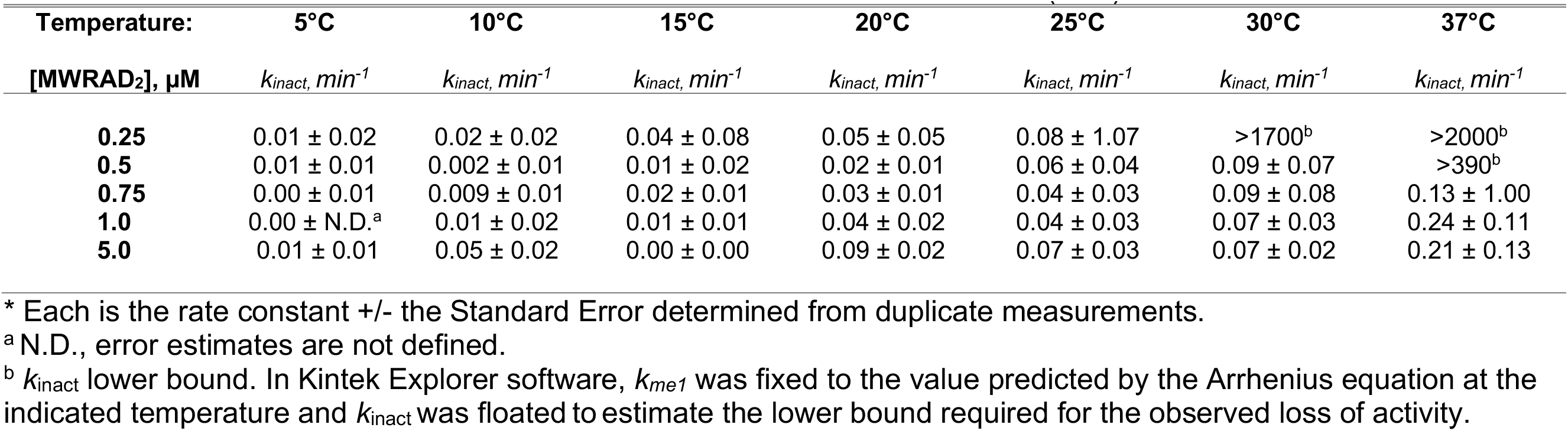
Pseudo-first order rate constants for MWRAD_2_ irreversible inactivation (*k*_*inact*_)

| Temperature: | 5°C | 10°C | 15°C | 20°C | 25°C | 30°C | 37°C |
| --- | --- | --- | --- | --- | --- | --- | --- |
| [MWRAD<sub>2</sub>], μM | $k_{inact}, \text{min}^{-1}$ | $k_{inact}, \text{min}^{-1}$ | $k_{inact}, \text{min}^{-1}$ | $k_{inact}, \text{min}^{-1}$ | $k_{inact}, \text{min}^{-1}$ | $k_{inact}, \text{min}^{-1}$ | $k_{inact}, \text{min}^{-1}$ |
| **0.25** | 0.01 ± 0.02 | 0.02 ± 0.02 | 0.04 ± 0.08 | 0.05 ± 0.05 | 0.08 ± 1.07 | >1700<sup>b</sup> | >2000<sup>b</sup> |
| **0.5** | 0.01 ± 0.01 | 0.002 ± 0.01 | 0.01 ± 0.02 | 0.02 ± 0.01 | 0.06 ± 0.04 | 0.09 ± 0.07 | >390<sup>b</sup> |
| **0.75** | 0.00 ± 0.01 | 0.009 ± 0.01 | 0.02 ± 0.01 | 0.03 ± 0.01 | 0.04 ± 0.03 | 0.09 ± 0.08 | 0.13 ± 1.00 |
| **1.0** | 0.00 ± N.D.<sup>a</sup> | 0.01 ± 0.02 | 0.01 ± 0.01 | 0.04 ± 0.02 | 0.04 ± 0.03 | 0.07 ± 0.03 | 0.24 ± 0.11 |
| **5.0** | 0.01 ± 0.01 | 0.05 ± 0.02 | 0.00 ± 0.00 | 0.09 ± 0.02 | 0.07 ± 0.03 | 0.07 ± 0.02 | 0.21 ± 0.13 |

\* Each is the rate constant +/- the Standard Error determined from duplicate measurements.
<sup>a</sup>N.D., error estimates are not defined.
<sup>b</sup> $k_{inact}$ lower bound. In Kintek Explorer software, $k_{me1}$ was fixed to the value predicted by the Arrhenius equation at the indicated temperature and $k_{inact}$ was floated to estimate the lower bound required for the observed loss of activity.

**Figure 8:**
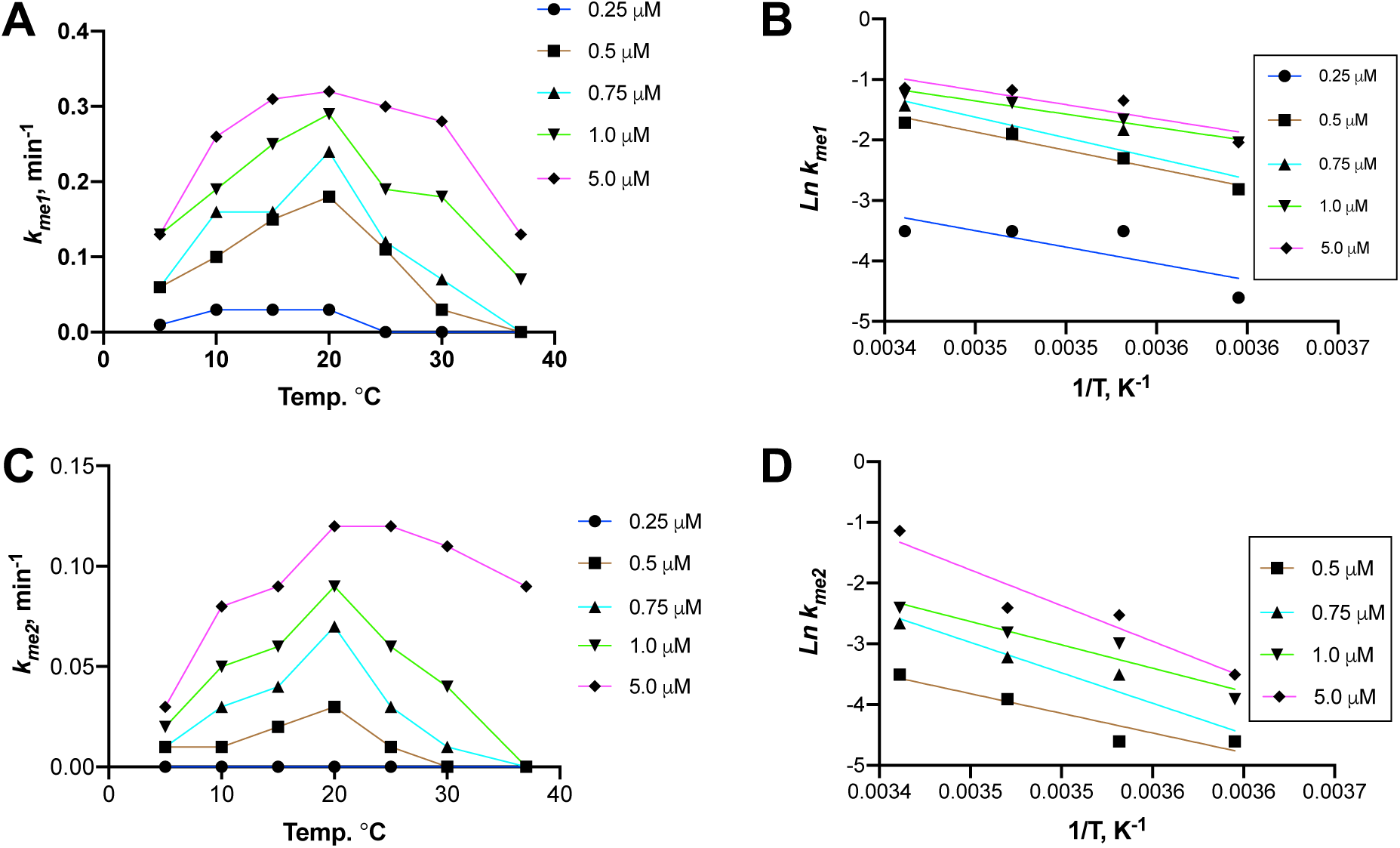
Effect of temperature on MWRAD_2_ enzymatic activity. (A and C), rates of H3K4 mono-(A) and dimethylation (C) plotted as a function of temperature. Arrhenius behavior (defined as a doubling of the rate for every 10°C increase in temperature) was observed between 5°C and 20°C for most concentrations. (B and D), Arrhenius plots for H3K4 mono-(B) and dimethylation (D) for the data collected between 5°C and 20°C. The lines represent linear regression fits to the data collected at the indicated MWRAD_2_ concentrations. *E_a_* values were obtained from the slope of the Arrhenius fits, where *slope* = -(*E*_*a*_/*R*) at each enzyme concentration.

The minimum enzyme concentration resulting in complete conversion into the mono- and then dimethylated forms was 1.0 μM at 15°C (Fig. 7). Slightly higher activity was observed at the same enzyme concentration at 20°C, but with evidence of significant enzyme inactivation resulting in failure to go to completion. Increased concentration extended the range of temperatures under which complete conversion could be observed. For example, at 5 μM enzyme concentration, complete conversion of the peptide into the dimethylated form was observed between 5°C and 30°C, with evidence of modest H3K4 trimethylation activity (7% - 15%) between 10°C and 25°C (Fig. 7). However, at 37°C, only ~25% of the peptide was converted into the dimethylated form before the enzyme was completely inactivated.

In general, the apparent pseudo first-order rate constants for mono- and dimethylation were correlated with the amount of holo-complex in the assay at every temperature between 5°C and 30°C, with Pearson correlation coefficients (r) ranging from 0.57-0.94 for monomethylation, and 0.46-0.86 for dimethylation. At 37°C, the correlation was less obvious due to the lack of detectable activity at the lowest concentrations. (r = 0.17 and 0.29 for mono and dimethylation, respectively). In contrast, the parameter that was most highly correlated with the amount of holo-complex present in the assay at all temperatures was the rate of irreversible enzyme inactivation (*k*_*inact*_), with Pearson r values ranging between −0.84 and −0.99, depending on the concentration tested. These results are consistent with the conclusions from the *s*_*w*_ isotherm analysis, in that holo-complex formation prevents individual subunits from sampling potential non-productive folding intermediates, some of which lead to irreversible enzyme inactivation. These results also raise questions about how cells manage to prevent loss of enzymatic activity at physiological temperatures.

### Induced high local concentration within a biomolecular condensate alters the assembly and enzymatic activity of the MLL1 core complex

Both hydrodynamic and enzymatic assays suggested that higher local concentrations of subunits would promote complex formation and enzymatic activity at physiological temperatures. However, given the low concentration of MLL1 in cells (which has been estimated to be femtomoles per mg of nuclear extract (65)), it is likely other factors are required to promote complex assembly. MLL1 has been shown to localize in discrete puncta in mammalian cell nuclei (66), raising the possibility that it could be regulated by induced high local concentration in liquid-liquid phase-separated (LLPS) particles, such as those found in transcription factories (67,68). Liquid-liquid phase separation has been shown to increase local protein concentration of proteins and ligands by up to 10,000-fold (69). Common features of proteins that undergo phase separation include primary sequences with regions of low complexity, or intrinsically disordered regions, that provide the numerous transient multivalent interactions required for liquid-liquid de-mixing (70). Indeed, examination of the primary sequence of MLL1 by IUPred (71) reveals that the majority of its sequence is predicted to be intrinsically disordered (Fig. 9A). In addition, the MLL1 construct used in this investigation and each WRAD_2_ subunit shows significant regions of predicted disorder (Fig. S2). To determine if the catalytic module of the MLL1 core complex may also be regulated by phase separation, we examined MWRAD_2_ using differential interference contrast (DIC) microscopy at concentrations up to 75 mg/ml but observed no evidence for phase separation (not shown). However, since a previous investigation showed increased enzymatic activity of the MLL1 core complex with reduced ionic strength (72), we tested whether reduced ionic strength may also regulate the LLPS properties of the MLL1 core complex.

**Figure 9:**
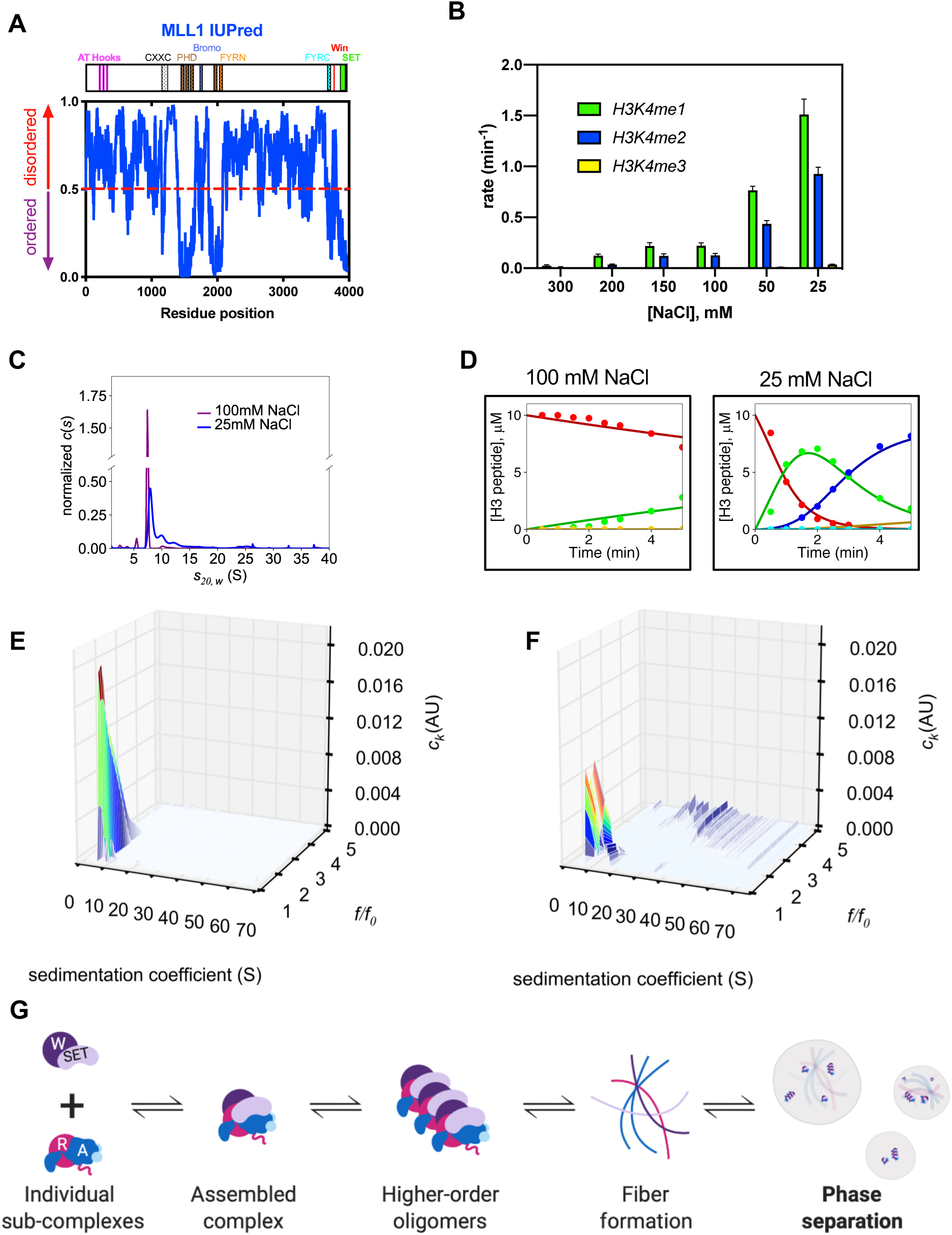
MLL1 core complex enzymatic activity is increased under conditions that induce phase separation. (A) IUPred disorder prediction (71) for the full-length MLL1 protein. Uniprot sub-domain boundaries are shown in the schematic above and are summarized in Table S6. (B) Comparison of 5 μM MLL1 core complex enzymatic activity at different ionic strengths at 25°C. (C) SV-AUC comparison of 5.0 μM MWRAD_2_ *c(s)* distributions at 100 mM (purple) and 25 mM (blue) NaCl. (D) Reaction time courses of 5.0 μM MWRAD_2_ at 100 mM NaCl (left panel) and 25 mM NaCl (right panel) at 25°C. Each time point represents the mean concentration of each peptide species, and solid lines show the fit using Scheme 6. Peptide species were H3K4me0 (red), H3K4me1 (green), H3K4me2 (blue), and H3K4me3 (yellow). (E and F) Size and shape analyses (*c(s,f*_*r*_)) from SV-AUC runs of 5.0 μM MWRAD_2_ in buffer with either 100 mM (E) or 25 mM (F) NaCl, each at 25°C. (G) A schematic of the build-up of higher-order oligomers and subsequent fiber formation preceding phase separation (Created with BioRender.com).

First, we compared MLL1 core complex activity at several different ionic strengths at 25°C using quantitative MALDI-TOF mass spectrometry. Consistent with the previous report, we found that the enzymatic activity was significantly increased in buffers with sub-physiological ionic strength (Fig. 9B, D). While there was relatively little difference in mono- or dimethylation activities between 200-100 mM NaCl, mono- and dimethylation activity was increased 15- and 12-fold, respectively, when the NaCl concentration was reduced from 100 to 25 mM (Fig. 9B, D). To better understand the reason for increased enzymatic activity at lower ionic strength, we compared the hydrodynamic properties of the 5 μM MWRAD_2_ complex at 100 mM and 25 mM NaCl at 25°C using SV-AUC (Fig. 9C). Strikingly, comparison of *c(s)* profiles showed hydrodynamic changes in the complex that resembled those of HP1α that was induced to undergo phase separation (73). The relatively monodisperse peak of the MLL1 core complex at physiological ionic strength (Fig. 9C, purple line) becomes more polydisperse when ionic strength is reduced (Fig 9C, blue line), with peaks at 8.1 *S* (54%), 10.0 *S* (21%) and 12.3 *S* (~10%), along with several higher molecular weight species that collectively account for ~16% of the total signal. We also noticed that the relative distribution among these species shifts to larger *S*-values in a concentration-dependent manner, which is more pronounced with even lower ionic strength (Fig. S3). These results suggest that the increased activity of the MLL1 core complex with lower ionic strength is associated with hydrodynamic alterations of the complex that could include conformational alterations, oligomerization, aggregation, and/or phase separation.

Because the standard *c(s)* analysis uses a single weight-average frictional coefficient of all particles to fit the data (74), the polydispersity of the sample at low ionic strength shown in Fig. 9C prevents accurate molecular weight estimates of each species – and thus our ability to distinguish among the different hypotheses. We therefore performed a two-dimensional size and shape distribution analysis *(c(s,f*_*r*_)) of the SV-AUC data, which allows estimation of the frictional coefficients and average molar masses of each species in a complex distribution (75). The *c(s,f*_*r*_) distribution of MWRAD_2_ at ~100mM NaCl showed a single peak with the typical experimental *s\**-value of the complex, but encompassing a fairly broad range of frictional ratios between 1.0 and 3.0, with a weight average frictional coefficient of ~1.5 (Fig. 9E). The estimated average molecular mass using this frictional coefficient and *S*-value was ~190 kDa, which is in fairly good agreement with the theoretical mass of the monomeric complex (205 kDa). In contrast, in low ionic strength buffer, the *c(s,f*_*r*_) distribution showed that the majority of the signal is divided among several peaks with larger *S*-values that ranged between 9 and 16, with evidence of several larger molecular weight species ranging between 20-70 *S* (Fig. 9F). Several of the peaks between 9 and 13 *S* had frictional ratios that range between 1.1-1.2, which gave mass estimates between 140-230 kDa. Because these species have relatively similar molar mass estimates, these *S*-values likely correspond to species with increasingly compact conformations of the monomeric MLL1 core complex. The peak at ~16 *S* gives a mass estimate of ~350 KDa, which is indicative of a reaction boundary between monomeric and dimeric complexes. These results suggest that lower ionic strength allows the complex to sample different conformational states, some of which are more compact, and some that allow oligomerization of the MLL1 core complex. Consistent with this interpretation, these larger *S*-value species become increasingly more populated in an MWRAD_2_ concentration-dependent manner (Fig, S3).

The *c(s,f*_*r*_) analysis also showed several discrete species with *S*-values between 20-70 *S* with a broad range of frictional ratios ranging between 3-5 (Fig. 9F). Integration of these peaks gave mass estimates starting at ~3.7 MDa, which approximates an 18-mer of MWRAD_2_, with each discrete species at higher *S*-values approximating the addition of one MWRAD_2_ dimer. This hydrodynamic behavior is indicative of fiber-like material (76) and could reflect various sizes of insoluble aggregates, or the fiber-like polymerization that is predicted to precede the formation of phase separated droplets (Fig. 9G) (70,77). To distinguish these hypotheses, we examined enzymatic reaction mixtures at 100mM or 25 mM NaCl using DIC microscopy. Surprisingly, despite using a relatively low concentration of enzyme (5 μM), the low ionic strength reaction mixture showed evidence of spherical LLPS droplets (Fig. 10B) that were absent in the 100mM NaCl reaction mixture (Fig. 10A). No visible evidence of protein precipitation was observed. The droplets were small and mobile, but did not appear to fuse, which is a common feature of particles induced to undergo LLPS (77). However, addition of a crowding agent (dextran; 7% w/v) to the reaction mixture resulted in LLPS droplets with larger diameters and observable fusion events that could be detected by DIC microscopy (Fig. 10C and movie S1). Importantly, the droplets disappeared in the presence of 5% 1,6 hexanediol (Fig. 9D), which has been shown to disrupt LLPS droplets formed by other proteins (78). We also note that in the presence of dextran, similar LLPS droplets (Fig. S4) and hydrodynamic behavior Fig. S5) could be observed at concentrations of NaCl that more closely approximated physiological ionic strength.

**Figure 10:**
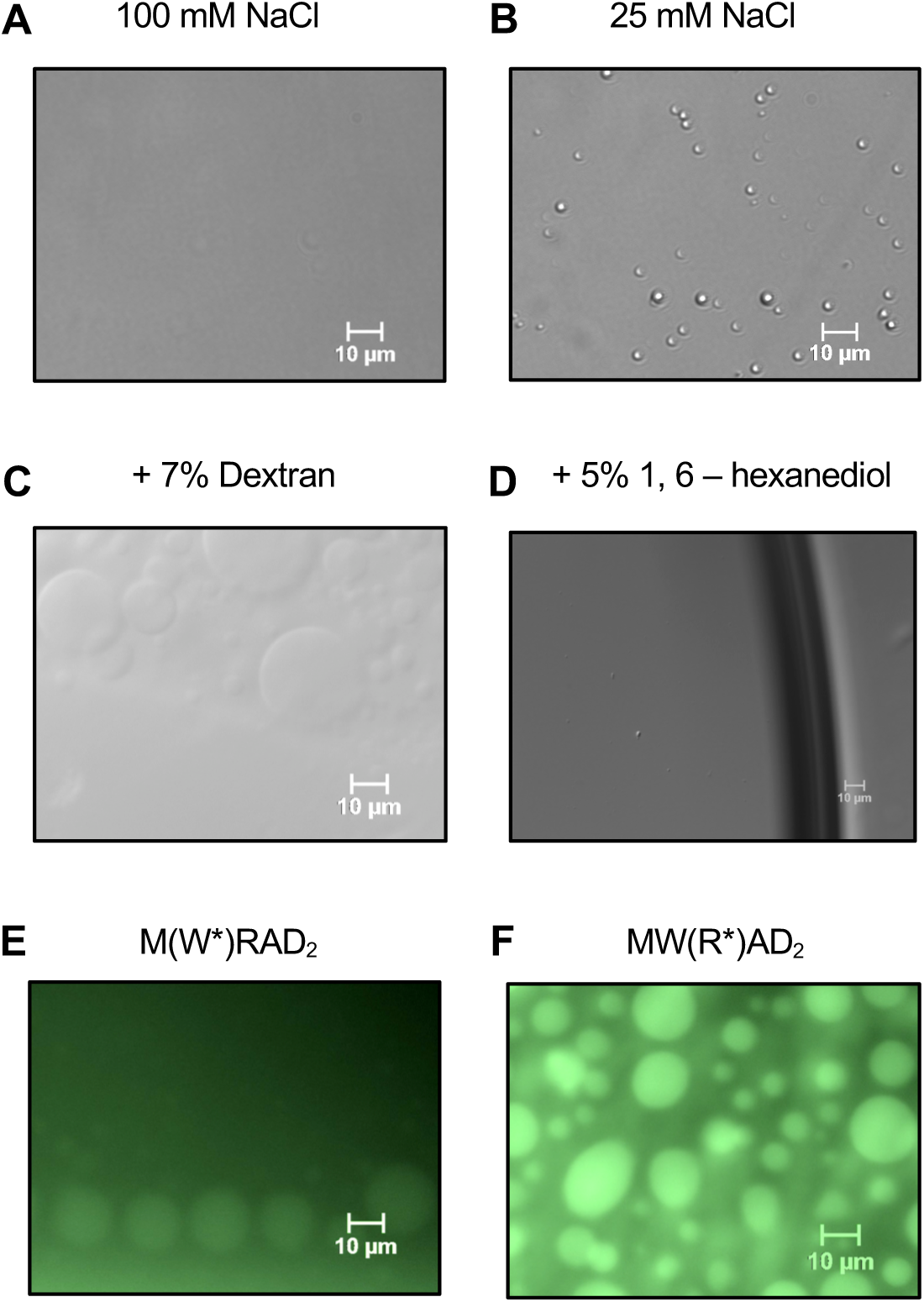
The MLL1 core complex phase separates at a low concentration and physiological ionic strength. (A,B) DIC microscopy images of MLL1 core complex enzymatic reactions at 100 mM (A) or 25 mM (B) NaCl. Each reaction contained 5.0 μM MWRAD_2_, 100 μM H3^1-20^ peptide and 250 μM AdoMet in reaction buffer at 25°C. (C) The same as in (B) but with 7% dextran (see also Supplementary movie S1). (D) Same as in (C) but with 5% 1,6 hexanediol. (E,F) Fluorescence microscopy images of the MLL1 core complex assembled with AlexaFluor 488-labeled WDR5 (E) or RbBP5 (F) subunits (see also supplementary movies S3 and S4). The conditions were 5.0 μM gel filtration-purified complex (see supplementary Fig.S7) in reaction buffer with 10 μM H3^1-20^ peptide, 250 μM AdoMet, and 150 mM NaCl.

Since we observed that higher concentrations of the histone H3 peptide alone showed evidence of phase separation (not shown), we next determined whether the LLPS droplets we observed contained only histone H3 or if they also contained the MLL1 core complex. To do this, we assembled the MLL1 core complex with fluorescently-labeled WDR5 (W*) or RbBP5 (R*) subunits and, after purification by SEC, we tested for their ability to phase separate using fluorescence microscopy. SEC elution profiles were similar to that of unlabeled complex (Fig S6A) and SDS-PAGE showed that each fluorescent subunit eluted in a stoichiometric complex with unlabeled subunits (Fig. S6B). In addition, control experiments with each complex showed that the fluorescent tag had minimal effect on enzymatic activity (Fig. S6C, D). When reactions were examined using fluorescent microscopy, both fluorescently labeled complexes were present in the buffer and inside the droplets (Fig. 10E, F and movies S2, S3). These results suggest that the catalytic module of the MLL1 core complex is in an equilibrium between phases both inside and outside of the LLPS droplets.

Lastly, to determine if LLPS formation rescues enzymatic activity at physiological temperature, we compared methylation kinetics of different concentrations of the MLL1 core complex among reaction mixtures containing 200 mM or 25 mM NaCl at 37°C. As described above, at near physiological ionic strength, none of the reactions went to completion, even after 24-hour incubation, mainly due to rapid enzyme inactivation at 37°C (Fig. 11, left column). In contrast, in low ionic strength buffer, most of the tested concentrations showed at least 80% conversion to the dimethylated form of H3K4 after only 5 minutes (Fig. 11, right column). At the highest concentrations tested (5 μM) the pseudo-first order rate constants for mono- and di-methylation increased 62- and 50-fold, respectively, with no evidence of enzyme inactivation (Table 5). Lastly, unlike the reactions using higher ionic strength, at low ionic strength, the reactions better approximated true single-turnover conditions with rates that were strictly dependent on enzyme concentration and not substrate concentration (79) (Fig. S7), as would be expected upon induced high-local concentration of enzyme within a biomolecular condensate.

**Table 5.**
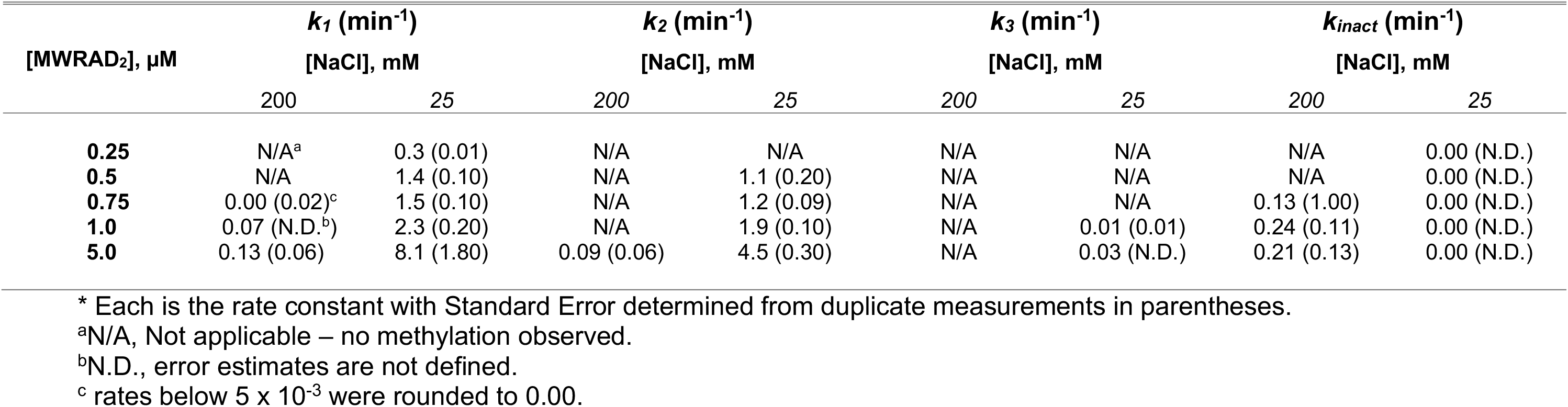
Pseudo-first order rate constants for 5 µM MWRAD_2_ at 37°C in high (200mM) and low (25mM) NaCl reaction buffer*

| [MWRAD<sub>2</sub>], $\mu\text{M}$ | *k*<sub>1</sub> (min<sup>-1</sup>) | | *k*<sub>2</sub> (min<sup>-1</sup>) | | *k*<sub>3</sub> (min<sup>-1</sup>) | | *k*<sub>inact</sub> (min<sup>-1</sup>) | |
| --- | --- | --- | --- | --- | --- | --- | --- | --- |
|  | [NaCl], mM |  | [NaCl], mM |  | [NaCl], mM |  | [NaCl], mM |  |
|  | 200 | 25 | 200 | 25 | 200 | 25 | 200 | 25 |
| **0.25** | N/A<sup>a</sup> | 0.3 (0.01) | N/A | N/A | N/A | N/A | N/A | 0.00 (N.D.) |
| **0.5** | N/A | 1.4 (0.10) | N/A | 1.1 (0.20) | N/A | N/A | N/A | 0.00 (N.D.) |
| **0.75** | 0.00 (0.02)<sup>c</sup> | 1.5 (0.10) | N/A | 1.2 (0.09) | N/A | N/A | 0.13 (1.00) | 0.00 (N.D.) |
| **1.0** | 0.07 (N.D.<sup>b</sup>) | 2.3 (0.20) | N/A | 1.9 (0.10) | N/A | 0.01 (0.01) | 0.24 (0.11) | 0.00 (N.D.) |
| **5.0** | 0.13 (0.06) | 8.1 (1.80) | 0.09 (0.06) | 4.5 (0.30) | N/A | 0.03 (N.D.) | 0.21 (0.13) | 0.00 (N.D.) |

\* Each is the rate constant with Standard Error determined from duplicate measurements in parentheses.
<sup>a</sup>N/A, Not applicable – no methylation observed.
<sup>b</sup>N.D., error estimates are not defined.
<sup>c</sup> rates below $5 \times 10^{-3}$ were rounded to 0.00.

**Figure 11:**
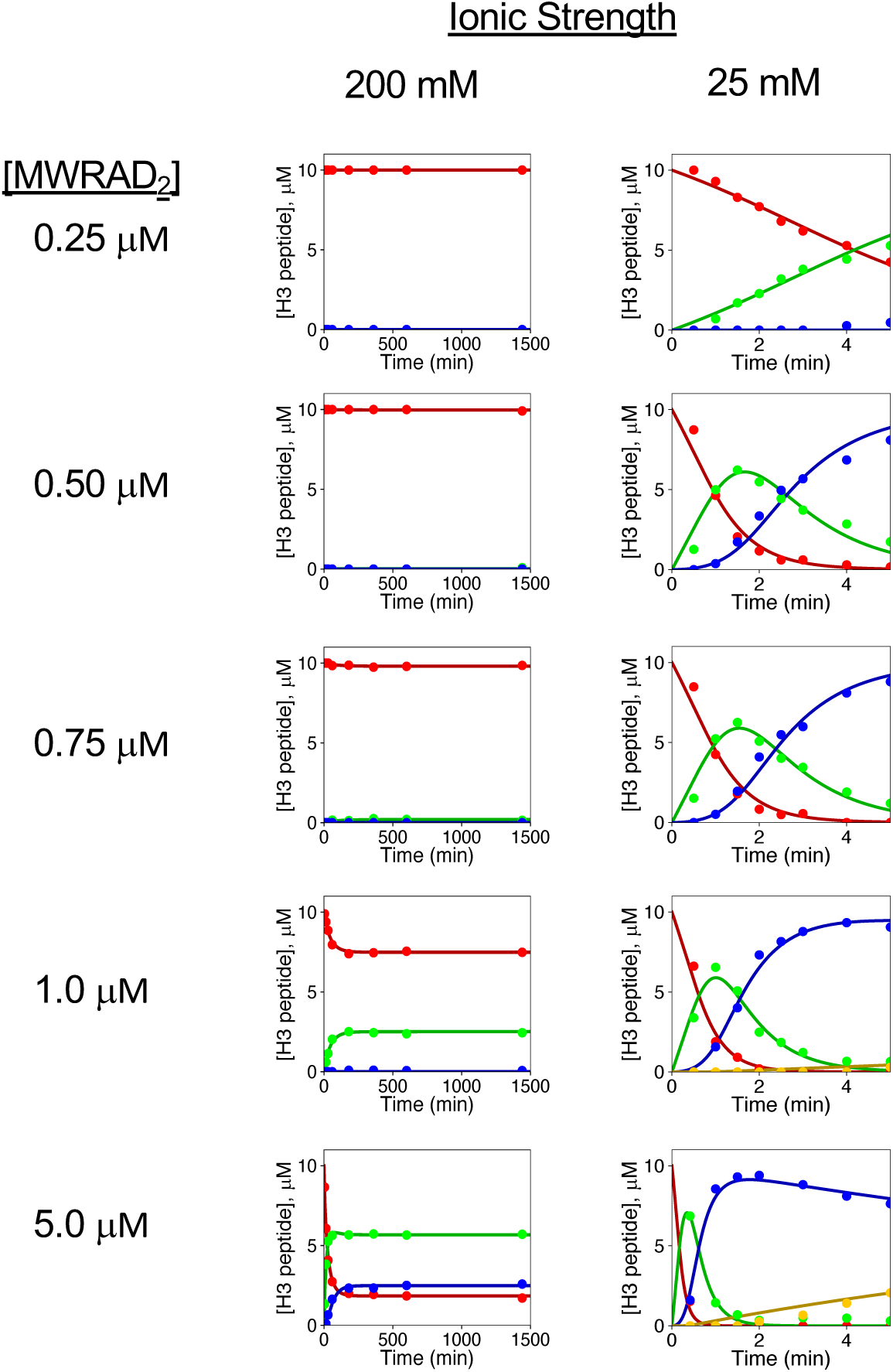
Enzymatic activity of the MLL1 core complex at physiological temperature under phase separation conditions. Comparison of MLL1 core complex enzymatic activity at the indicated concentrations at 37°C in high (200 mM NaCl) vs. low (25 mM NaCl) ionic strength reaction buffers. The 200mM NaCl panels (left) from Fig. 7 are shown again here for the purpose of comparison. Each time point represents the mean concentration of each peptide species and solid lines show the fit using Scheme 6 (Fig.6) or a modified form of Scheme 6 to account for trimethylation. The resulting pseudo-first order rate constants are summarized in Table 5. Peptide species were H3K4me0 (red), H3K4me1 (green), H3K4me2 (blue), and H3K4me3 (yellow). Note the time scale differences required for the high vs. low ionic strength reactions.

All together, these results are consistent with the hypothesis that induced high-local concentration within a biomolecular condensate overcomes the thermodynamic barrier for MLL1 core complex assembly at physiological temperatures.

## DISCUSSION

Numerous studies have established the role of MLL1 in the regulation of the degree of H3K4 methylation in mammalian gene expression and human disease. While it has been shown that the SET domain has intrinsic H3K4 monomethylation activity, several studies have shown that multiple methylation depends on interaction of MLL1 with the WRAD_2_ sub-complex. While the molecular details for this product specificity switch are still in question, the idea that regulated complex assembly controls the spatial and temporal deposition of different H3K4 methylation states has significant experimental support. The importance of understanding the molecular details of this mode of regulation is demonstrated by studies showing targeted inhibition of the Win motif-WDR5 protein-protein interaction within the MLL1 core complex selectively reduces proliferation of MLL1-translocation leukemias and other cancer cells (42–44,80). These results suggest that molecules mimicking the Win motif, collectively called Win motif inhibitors, may be useful alternative or complementary therapeutics for cancer.

However, progress in exploiting this potential has been impeded by the lack of understanding of the biophysical and thermodynamic mechanisms that underlie MLL1 core complex assembly. The lack of standardized *in vitro* assay conditions has resulted in different conclusions regarding the mechanisms of multiple lysine methylation by SET1 family complexes and identification of the best inhibitors. For example, we previously found that the same Win motif inhibitor gives IC_50_ values that vary by more than an order of magnitude when assayed over a relatively narrow concentration range of the MLL1 core complex (0.5-1.8 µM) (81), suggesting complex assembly is relatively labile. Missing is a complete understanding of the conditions under which the complex is assembled when assayed *in vitro*. This is crucial not only for our ability to compare the potency and specificity of different inhibitors, but also for establishing a baseline for understanding how the dynamics of MLL1 core complex assembly is regulated in cells.

In this investigation, we systematically characterized the hydrodynamic and kinetic properties of a reconstituted human MLL1 core complex under a variety of assay conditions. As expected, we found that complex assembly is highly concentration and temperature dependent. Consistent with the hypothesized hierarchical assembly pathway, we found that the holo-complex assembles through interactions between the MW and RAD_2_ sub-complexes, and that this assembly correlated with enzymatic activity. However, unexpectedly, we also found that the disassembled state of the complex is favored at physiological temperatures and at the sub-micromolar enzyme concentrations typically used in steady-state enzymatic assays (in which the substrate is in vast excess compared to the concentration of enzyme). We found that the complex disassembly results in rapid and irreversible enzyme inactivation under these conditions, likely because one or more subunits samples unproductive conformational states. Consistent with this conclusion, it was previously shown that overexpression of C-terminal fragments from the human SETd1A protein in mammalian cells depletes WRAD_2_ subunits from the endogenous SETd1A and SETd1B paralogs, resulting in their degradation (82). It is possible that in the cell, unproductive folding intermediates are limited by interaction with chaperones. Consistent with this hypothesis, HSP70 and HSP90 proteins have been found to co-purify with MLL1 super-complexes (31,33). In addition, HSP90 has been shown to be required for the stability of human MLL1 and the *Drosophila melanogaster* ortholog, *Trithorax*, which is important for homeotic gene expression (83). It remains to be determined if these or other chaperones interact with and regulate folding of the subunits of the catalytic module.

Our data suggest that the MW and RAD_2_ sub-complexes interact with a 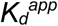 of ~6 μM at 37°C, raising the question of how complex forms in cells that contain relatively few molecules of MLL1, which has been estimated to be femtomoles per mg of nuclear extract (65). WRAD_2_ subunits appear to be present in cells in vast excess compared to that of MLL1 (65), which could help overcome the thermodynamic barrier to complex assembly. However, our previous demonstration that a stoichiometric excess of WDR5 inhibits the enzymatic activity of MLL3 (84) and MLL1 (unpublished) core complexes argues against this possibility. Our data suggest that inhibition by excess WDR5 results from saturation of available binding sites on the RAD_2_ sub-complex, which would prevent its interaction with the MW sub-complex. These results suggest that cellular pools of WDR5 may need to be compartmentalized to prevent this form of inhibition of SET1 family complexes. This may explain why WDR5 over-expression is associated with several poor outcome malignancies, including bladder, breast, colon, and prostate cancers, leukemias and hepatocellular carcinomas (80,85–89).

Alternative possibilities to overcome the barrier to complex formation in cells include interaction with other unknown proteins, cofactors, nucleic acids, post translational modifications, and/or by inducing a high local concentration of MWRAD_2_ subunits within a phase separated compartment. While there is evidence that phosphorylation and long non-coding RNAs regulate the function of MLL family complexes (90,91), it is currently unclear if these mechanisms would overcome the barrier to MLL1 core complex assembly at physiological temperatures. Our data suggests that the barrier to complex formation is overcome in cells by concentration of subunits in biomolecular condensates, such as those found in transcription factories (67). Biomolecular condensates are membraneless liquid-like organelles, or intracellular phase-separated compartments, that function to concentrate proteins and nucleic acids to regulate a variety of biological processes (77,92). This form of compartmentalization has been shown to have variable effects on the activity of enzymes, ranging from a 2-70-fold stimulation in the rate of enzyme or ribozyme-catalyzed cleavage reactions, to inhibition of catalyzed reactions, protein conformational alterations and increased thermal resistance (79,93–97). While there are a number of recent examples of chromatin and chromatin-associated proteins that undergo LLPS in mechanisms that may regulate heterochromatic gene silencing (69,73,98–100), to our knowledge, there is currently no evidence demonstrating LLPS regulation of enzymatic activity of a histone modification enzyme.

Our data suggest that concentration of the MLL1 core complex in a biomolecular condensate overcomes the barrier to complex assembly at physiological temperatures, resulting in histone methyltransferase activity that is increased by at least 30-60-fold (Table 5), depending on the enzyme concentration in the assays. However, the molecular mechanism for how compartmentalization stimulates MWRAD_2_ activity is likely more complex. This is because the hydrodynamic properties of the complex change under phase separation conditions, likely involving conformational changes and oligomerization that may be prerequisites for the multivalent interactions required for LLPS. It is interesting to note that in the absence of a crowding agent, these hydrodynamic changes begin to occur at the lower boundary of physiological ionic strength. This suggests a plausible regulatory mechanism in which small changes in ionic strength, possibly through compartmentalization, could have a large impact on MLL1 core complex activity. However, we also note that further lowering the ionic strength of the buffer (< 50 mM) resulted in detection of up to six methylation events on the same peptide (Fig. S8), suggesting reduced enzyme specificity. This result, may help explain contradictory results from different labs using different assay conditions. In addition, differences in the stability of MLL family complexes may underlie different conclusions about their relative activities. For example, we and others have observed that the MLL3 core complex is significantly more stable than the other MLL family complexes (53,84,101), which may account for observations suggesting that the MLL3 core complex is more active (72,102). However, we have found that when comparing enzymes under conditions where complexes are at least 80% assembled, there is little difference in the overall rate of H3K4 monomethylation among SET1 family complexes (84). Our results here underscore the importance of assaying enzymes under conditions where complexes are fully assembled, which in several cases may preclude the use of low enzyme concentrations typically used in steady-state kinetics studies.

Our results suggest a model in which MLL1 enzymatic activity is regulated in the cell at the level of complex assembly within a phase-separated transcription factory. Several lines of experimental evidence are consistent with this hypothesis. Early confocal microscopy studies showed that transcription occurs in a defined number of discrete sites within the cell called transcription factories (103,104), each containing a protein-rich core that encompasses RNA polymerase (Pol) II, co-activators, chromatin remodelers, transcription factors, histone modification enzymes, ribonucleoproteins, RNA helicases, splicing and processing factors (105). Indeed, peptides derived from WDR5 and DPY30, the two most abundant MLL1 core complex subunits (65), were found in purified RNA Pol II transcription factories (105). A phase separation model may explain, in part, immunofluorescence experiments showing that MLL1 has a punctate distribution within mammalian cell nuclei (66), which is a common feature of proteins that undergo LLPS (77). Furthermore, use of the PScore (106) and CatGRANULE (107) LLPS prediction programs show that MLL1, as well as all human MLL family proteins, have high phase separation probabilities (Table S3), as does Ash2L and Ash2L-containing sub-complexes (Table S4). In addition, it was recently demonstrated that the multivalent interactions provided by the carboxyl-terminal domain (CTD) of RNA polymerase (Pol) II are sufficient for formation of RNA Pol II LLPS clusters (78). Since several studies suggest that RNA Pol II interacts directly with MLL1 (108,109), it is possible they function together within phase-separated transcription factories. Consistent with this model, ChIP studies show that MLL1 and RNA Pol II co-localize at nucleosomes throughout the promoters and open reading frames of actively-transcribed genes (109). However, a puzzling aspect of this model is that, despite a study showing that MLL1 can be pulled-down from nuclear extracts with a recombinant GST-CTD fusion protein (109), Pol II appears to be absent in purified MLL1 super-complexes (30–33). It may be that co-localization within the same transcription factory is required for the interaction.

Combining our results on the assembly of the catalytic module with the observation that it follows a large region of predicted intrinsic disorder in the primary sequence of MLL1 (Fig. 9A), we propose a “swinging domain” model for the mechanism of action of the MLL1 core complex within cellular transcription factories (Fig. 12). A swinging domain is a common feature of enzyme complexes involved in multistep assembly pathways and are characterized by a structured mobile domain tethered to other components by conformationally flexible linker regions (110). This may explain why the low complexity region is conserved not only among MLL1 orthologs, but also in the primary sequences in all human SET1 family members, with the main differences being the length of the linker regions that precedes the SET domain (Fig. S9). This observation suggests that a swinging domain may be a conserved feature of SET1 family complexes (Fig. S10D) and linker length differences could be a unique regulatory feature that limits the range of nucleosomes that can be reached within different transcriptional compartments. This hypothesis deserves further investigation.

**Figure 12:**
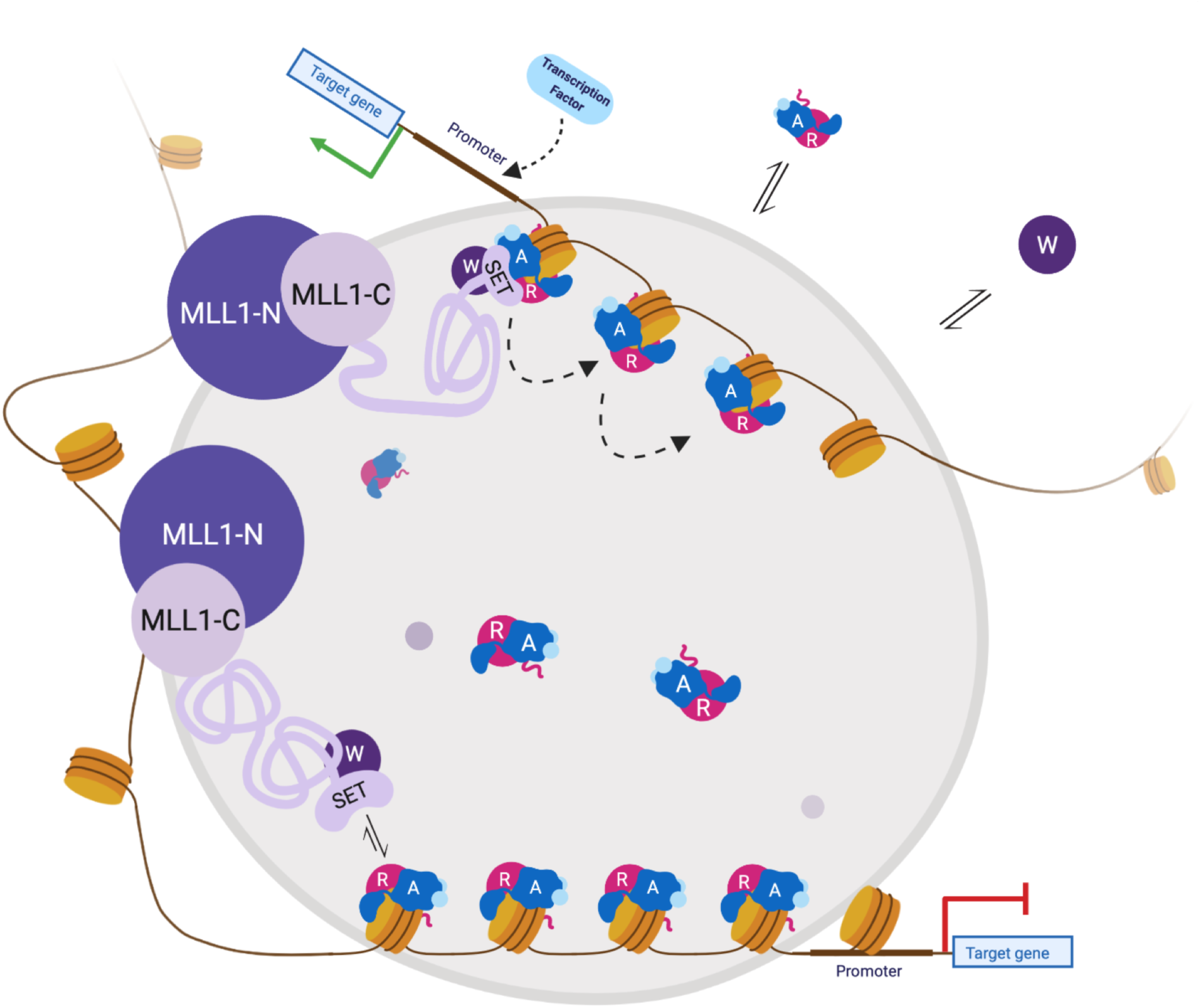
Swinging Domain Model for regulation of MLL1 core complex assembly and enzymatic activity within a transcription factory. MLL1 N-terminal region (MLL1-N) binds to DNA in or near a transcription factory using its DNA and chromatin-recognition domains. The C-terminal region (MLL1-C), which contains the SET domain, binds to WDR5 (W) to create the MW sub-complex, either before or after entry into the factory. The RbBP5, Ash2L, DPY-30 (RAD_2_) sub-complex binds nucleosomes. MW then uses a “swinging domain” mechanism within the phase condensate to move the SET domain-WDR5 around the transcription factory, where the high local concentration forces the assembly of the full MLL1 core complex and allows for H3K4 methylation of nucleosomes within the factory that have RAD_2_ already bound. This can be repeated multiple times within the factory, resulting in extensive H3K4 methylation of nucleosomes that go into the condensate. This methylation results in removal of nucleosomes and recruitment of transcription factors that, in turn, recruit RNA Polymerase II for transcription initiation. Once the chromatin leaves the factory, the reduction in local concentration results in the loss of the RAD_2_ sub-complex, as well as a high kinetic barrier to reassembly of MWRAD_2_, preventing any ectopic methylation. This figure was created with BioRender.com

A swinging domain model where the SET domain-WDR5 complex swings to different nucleosomes provides a satisfying explanation for how the relatively few molecules of MLL1 in the cell could methylate multiple nucleosomes in the promoter and open reading frames of genes as they move through the transcription factory (Fig. 12). This model also provides a plausible explanation for the observation of MLL1 and RNA Pol II co-localization in ChIP experiments without the necessity of a physical interaction. Given that RAD_2_ subunits are relatively abundant in cells and that the RAD_2_ sub-complex interacts with nucleosomes in the absence of the MW sub-complex (manuscript in preparation), concentration of both sub-complexes within a transcription factory could provide the energy required to overcome the barrier for holo-complex formation at physiological temperatures, resulting in activation of the histone methyltransferase activity of the MLL1 core complex. This model provides an elegant “switch-like” mechanism for spatiotemporal control of H3K4 methylation through the rapid formation or dissolution of biomolecular condensates, which would ultimately regulate the hierarchical assembly of the MLL1 core complex.

## ACKNOWLEDGEMENTS

We thank John Sfakis for help with the Fluorescent labeling experiments and Connie Mitra for assistance with low-salt MALDI-TOF activity assays. We thank Patty Kane for access to the Zeiss light microscope and Brian Haarer for assistance with the microscopy. We also thank Steve Hanes, Bruce Knutson and Alaji Bah for helpful suggestions and Ashley Canning and Michael Connelly for critical reading of the manuscript. This work was supported in part by R01C140522 (to M.S.C.).

## METHODS

### Protein Expression and Purification

Each of the human genes for the MLL1 SET domain (a.a. 3745-3969 – Uniprot #: Q03164), WDR5 (2-334 – P61964), RbBP5 (1-538 – Q15291) and Ash2L (1-534 – Q9UBL3-3) (111) were cloned into the pST44 polycistronic vector (112). The WDR5 subunit was cloned with an N-terminal 6x-Histidine tag followed by a Tobacco Etch Virus (TEV) protease cleavage site. Plasmids were transformed into Rosetta pLysS BL21 *E. coli* cells and plated on LB agar supplemented with 50 μg/mL carbenicillin and 20 μg/mL chloramphenicol (both from Gold Biotechnology). Individual colonies were used to inoculate a seed culture of 50mL of Terrific Broth II (MP Biomedicals), again supplemented with carbenicillin and chloramphenicol and grown overnight at 30°C. 20mL of the seed culture were used to inoculate 1L of Terrific Broth II media in baffled 2800mL flasks, maintaining the antibiotic resistance. Cultures were then grown for 2-4hrs at 37°C and 200RPM shaking until the O.D_.600_ reached ~1. Cultures were then chilled for 1hr at 4°C followed by induction with 1mM Isopropyl β-D-1-thiogalactopyranoside (IPTG – Gold Biotechnology), after which cells were grown for an additional 20-22hrs at 16°C with constant shaking. Cells were harvested by centrifugation at 4°C and pellets were flash frozen in liquid nitrogen and stored at −80°C until they could be lysed. Frozen cells were thawed and resuspended in 50 mL of lysis buffer (50mM Tris-HCl, pH 7.5; 300mM NaCl; 30mM Imidazole; 3mM dithiothreitol (DTT) and 1μM ZnCl_2_, supplemented with one tablet of EDTA-free protease inhibitor cocktail (Roche)), lysed with a microfluidizer, and cleared by centrifugation at 17,000RPM at 4°C for 30min. The supernatant was diluted to 250 mL in Buffer 1 (50mM Tris-HCl, pH 7.5; 300mM NaCl; 30mM Imidazole; 3mM DTT and 1μM ZnCl_2_) and flowed over a HisTrap 5mL nickel affinity column (GE) using an AKTA Purifier FPLC (GE) at a rate of 0.5 mL/minute. Bound complex was washed with 10 column volumes (CV) of Buffer 1 at 1 mL/min., and then eluted with a 25-CV linear gradient of Buffer 2 (Buffer 1 with 500 mM imidazole). Fractions containing the MWRA complex were pooled, supplemented with GST-6x-His-TEV protease to a final concentration of 0.1 mg/mL and dialyzed against Buffer 1 with three changes. The complex was then passed over a re-equilibrated HisTrap column and fractions from the flow-through containing the cleaved MWRA sample were collected, concentrated by ultrafiltration using a 30 kDa cutoff membrane to ~12 mg/mL, and further purified by size-exclusion chromatography (SEC) using a Superdex 200 (16/60) column (GE) pre-equilibrated with Buffer 3 (20mM Tris-HCl, pH 7.5; 300mM NaCl; 1mM TCEP and 1μM ZnCl_2_). A two-fold Molar excess of Human DPY-30 (1-99 – Q9C005), expressed and purified as previously described (36), was added to the MWRA sample and the resultant complex was purified with multiple rounds of SEC in buffer 3. Fractions containing purified MWRAD_2_ were concentrated to 12 mg/mL, aliquoted, flash frozen, and stored at −80°C until use. Individual subunits for Bayesian experiments were purified as previously described (36).

### Sedimentation Velocity-Analytical Ultracentrifugation

#### Experimental Procedures

All stock protein samples were thawed on ice, diluted to the desired concentration, and spun at 15,000RPM for 15min. at 4°C using a Thermo Scientific tabletop refrigerated centrifuge to remove any debris. Protein concentrations were measured with a NanoDrop spectrophotometer using the extinction coefficient *ε*^280^ of 248,954 M^−1^ cm^−1^, which was predicted from the amino acid sequence using ProtParam (113). 100 or 400 μL of diluted protein samples were then loaded into AUC cells containing 3- or 12-mm two-sector charcoal-Epon centerpieces (SpinAnalytical) assembled with quartz or sapphire windows. Matching buffer was loaded into the reference sector of each cell. AUC cells were then loaded into a Ti-60 4-hole Beckman-Coulter rotor, pre-equilibrated to the specific run temperature for at least 4hrs. Rotors were then inserted into the chamber of the centrifuge and allowed to re-equilibrate to experimental temperature for a minimum of 2hrs before initiation of the run. Sedimentation velocity analytical ultracentrifugation (SV-AUC) was performed using a Beckman-Coulter Proteomelab XL-A analytical ultracentrifuge equipped with absorbance optics. Each run was preceded by a 3000-rpm wavelength scan to detect cell leakage and to select the appropriate wavelength to ensure a starting absorbance of between 0.25 and 1.2 OD units. Wavelengths at or near the maximal absorbance for aromatics of 280 nm or peptide backbone of 230 nm were selected, depending on the protein concentration and pathlength of the centerpiece. Without slowing the rotor, a method scan of 50,000-rpm was initiated, and 200 scans/cell were collected with the time interval between scans set to zero. Each experiment was replicated in duplicate or triplicate.

#### Data Analysis

Lamm equation modeling of all SV-AUC results was performed using the continuous distribution (*c(s)*) method in SEDFIT (56). Maximum entropy (ME) regularization using a confidence level of P = 0.68 was performed to identify the most parsimonious distribution consistent with the data, and the fits for each experiment gave acceptable RMSD values ranging between 0.003 and 0.01. Density, viscosity and partial specific volume values were estimated by inputting the temperature, buffer reagents, and amino acid sequences of all five complex components (assuming a DPY-30 dimer) into the SEDNTERP program (114), and the values used are listed in Table S5. The resulting *c(s)* distributions were displayed and further analyzed using GUSSI (115). To determine the amount of holo-complex under each condition, distributions were integrated between *S*-values 6.8 and 7.6, which represents one standard deviation from the mean *S*-value of the holo-complex peak over all conditions, which was 7.2 +/− 0.4. For binding analyses, *c(s)* distributions were integrated from 0.5 to 9.5 *S* to derive the corresponding signal-weighted average sedimentation coefficients (*s*_*w*_), which were plotted as a function of loading concentration at each temperature and fit with mass action law models using the program SEDPHAT (116).

For Bayesian analyses of c(s) distributions, expected sedimentation coefficients were derived from separate SV-AUC experiments of individual subunits or assembled sub-complexes, which were each run at concentrations ranging from 0.25 to 5 μM at 25°C (the data for 0.25 μM runs are shown in Fig. S1). These values were then used in ME regularization as prior expectation restraints to give *c*^(*p*)^(*s*) distributions of the holo-complex at 25°C. Prior expectations for sub-complexes or individual subunits were implemented as Gaussians in SEDFIT for Bayesian analysis, with a peak width of sigma = 0.2 *S* and centered at the weight-average *S*-value of the main peak observed in the individual experiments with an amplitude of 0.05 OD units. Since the prior expected *S*-values for WDR5 or RbBP5 overlapped when run in individual experiments, they were used as prior expectations in *c*^(*p*)^(*s*) distributions to test the concerted assembly mechanism with the same weight average *S*-value but with an amplitude that was doubled (Fig. 5D). Each *c*^(*p*)^(*s*) distribution was fit with the same prior expectation for MWRAD_2_, which used the weight-average *S*-value determined at 25°C and 0.25 μM with a width of sigma = 0.4 *S* and an amplitude of 0.3 OD units.

For the *c(s,f*_*r*_) analysis, the data were first imported into SEDFIT with reduced radial resolution (0.006cm compared to the default 0.003cm) and loading every second scan, to reduce the computational power required (60). These were fit using the *c(s, f*_*r*_) method in SEDFIT with resolutions of 50 for both the sedimentation coefficient and frictional ratio dimensions.

### Methyltransferase Activity Assay

MWRAD_2_ complex was assayed using a label-free quantitative MALDI-TOF mass spectrometry assay (36). Each 20 μL reaction consisted of varying concentrations of MWRAD_2_, 250 μM S-adenosylmethionine (AdoMet) and reaction buffer (50 mM Tris, pH 9.0; 200 mM NaCl; 5% (v/v) glycerol; 1 μM ZnCl_2_; 3 mM DTT), which were preincubated for 5 minutes at the experimental temperature in a thermocycler. Reactions were initiated by the addition of temperature-pre-equilibrated histone H3 peptide (residues 1-20, with an additional C-terminal GGK-biotin moiety) to a final concentration of 10 μM. At various timepoints, a 2 μL aliquot was removed and quenched by mixing with 2 μL of 1% trifluoroacetic acid (TFA). Quenched reactions were stored at −20°C until they could be analyzed. Upon analysis, samples were thawed and 1 μL of each was mixed with 4 μL of α-cyano-4-hydroxycinnamic acid in 0.05% TFA and 50% acetonitrile. 2 μL of this mixture for each time point was spotted onto a ground steel target plate and allowed to dry at room temperature for 3-12 hours. Spectra were acquired on a Bruker Autoflex III MALDI-TOF mass spectrometer in reflectron mode. Each spectrum was the sum of at least 1000 individual laser shots, obtained from five different positions around the spot, with 200 shots at each position. Using FlexAnalysis software (Bruker), the intensities of the unmodified (m/z 2651 Da), mono-(m/z 2665 Da), di-(m/z 2679 Da), and trimethylated (m/z 2693 Da) species were summed to obtain the total intensity. The relative amount of each species was then determined by dividing the intensity of each methylation state by the total intensity at each time point and multiplied by the starting substrate concentration (10μM) to give the micromolar concentration of each methylation state. These data were then plotted as a function of time for kinetics analyses.

Fitting of the data was performed using the numerical integration of rate equations approach implemented in KinTek Explorer software version 6.3 (61). For reaction schemes incorporating the complex dissociation step, the ratio (*k*_*off*_/*k*_*on*_) was constrained to be equal to estimated 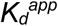 for complex dissociation at each temperature determined from the sedimentation velocity *s_w_* isotherm analysis, with the *k_on_* fixed at the limit of diffusion. All other non-variable parameters were fixed with non-rate limiting values. Confidence contour analysis using a *Chi*^*2*^ threshold of 0.9 was used to obtain estimates for the extent to which each variable parameter was constrained by the data.

### Labeling and assembly of fluorescent MWRAD_2_ complexes

Recombinant WDR5 or RbBP5 were expressed and purified as previously described (37). Purified proteins at ~14 mg/ml were dialyzed into labeling buffer composed of 20 mM HEPES, pH 7.0; 300 mM NaCl; 1 mM TCEP and 1 μM ZnCl_2_. The neutral pH was chosen to facilitate selective labeling of the free amino terminus of the protein, which has a lower *pK*_*a*_ than the primary amines of the lysine side chains (117). The protein was mixed with AlexaFluor™ 488 NHS Ester (Invitrogen) in a 1:6 (for WDR5) or 1:5 (for RbBP5) molar excess of label and reacted for 3 hours at 4°C. The entire reaction volume for each protein was then loaded onto a Superdex™ 200 10/300 GL size-exclusion column (GE) to separate the labeled protein from the unreacted fluorophore. The labeled protein fractions were then combined and concentrated by ultrafiltration in a 10,000 MWCO concentrator (Millipore). Once concentrated, the degree of labeling was determined using the equations shown below:

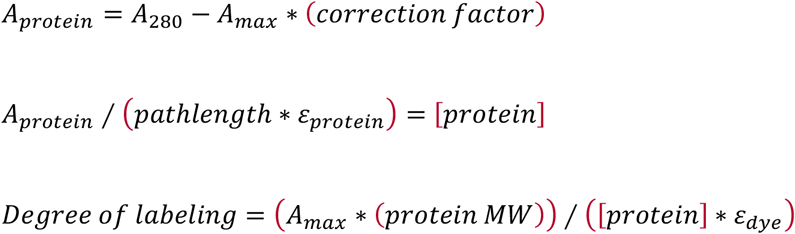

The degree of labeling for WDR5 (W*), was found to be 1.1, or ~ 1 molecule of fluorophore for each molecule of WDR5. The degree of labeling determined for RbBP5 (R*) was 1.9, or ~2 molecules of fluorophore per molecule of RbBP5. Each labeled protein was then mixed in equivalent molar ratios with the other recombinant, unlabeled complex components and loaded onto a Superdex™ 200 10/300 GL size-exclusion column and fractions containing stoichiometric complex were pooled, concentrated, and stored at −80°C until use.

### Liquid-Liquid Phase Separation (LLPS) Assays

MWRAD_2_ at a concentration of 5 μM was mixed with H3^1-20^ peptide (100 – 500 μM) and 250 μM SAM in either physiological (~100-150 mM) or sub-physiological (~25-50 mM) NaCl buffers containing (50 mM Tris, pH 9.0, 1 μM ZnCl_2_, 3 mM DTT and 5% (w/v) glycerol) in the presence or absence of 7% (w/v) Dextran Sulfate (avg. M.W. = 500,000 Da) as a crowding agent. 1 μL of each sample was pipetted into the depression of 12-well precleaned frosted end Bioworld microscope slide, covered by a cover slip, and observed on a Zeiss light microscope in DIC mode at 40x magnification. Single images and movies were taken using a Hamamatsu camera connected to the microscope. All images taken are of samples at room temperature (~23°C). In addition to DIC, M(W*)RAD_2_ or MW(R*)AD_2_ were imaged with the FITC filter activated. As a control for phase separation, reaction mixtures were compared in the presence and absence of 5%1,6 hexanediol.

