## Supplementary Figures and Tables for "Hierarchical assembly of the MLL1 core complex within a biomolecular condensate regulates H3K4 methylation"

#### Figure S1 – Namitz, Tan and Cosgrove

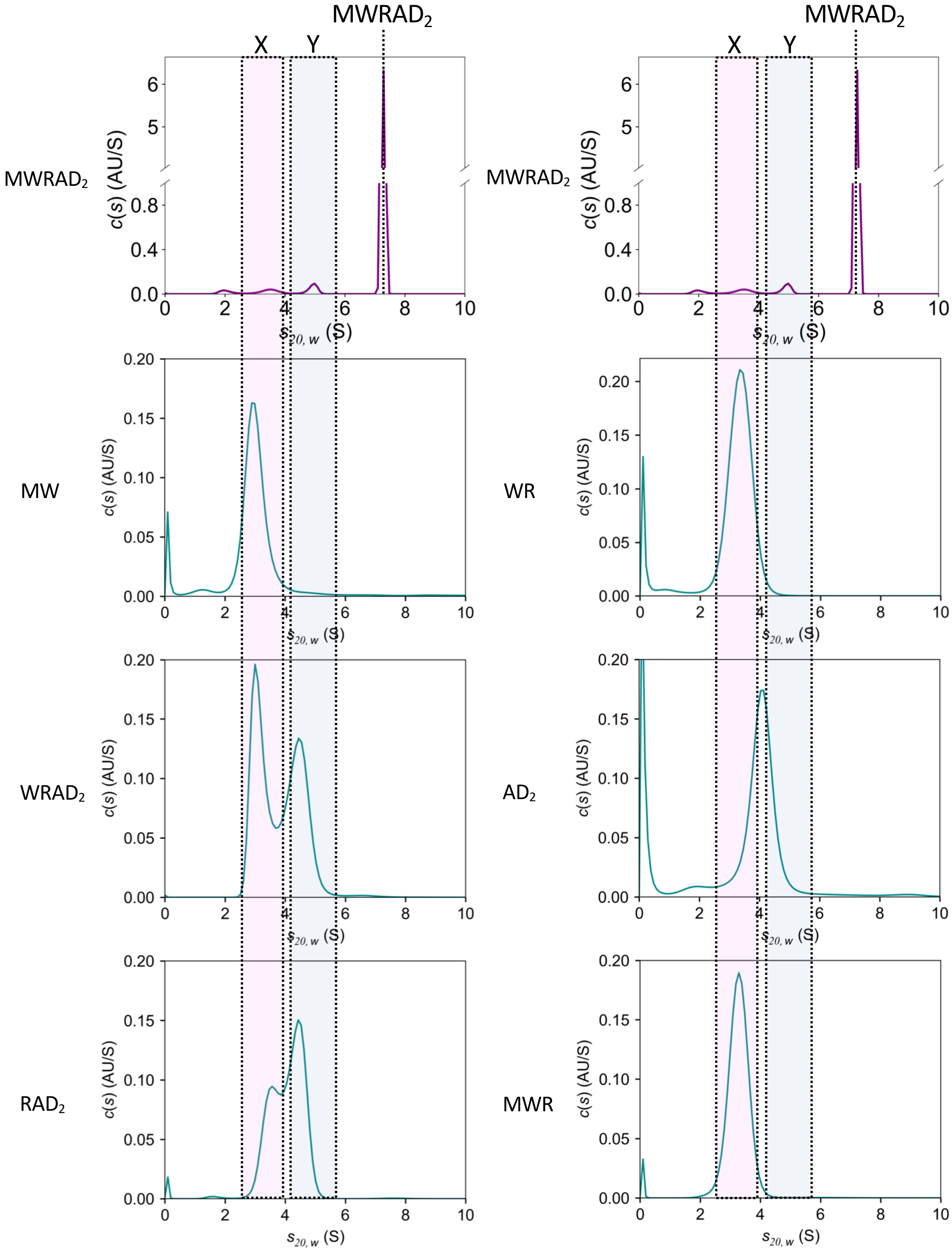


**Figure S1: Determination of expected sub-complex *S* values for Bayesian analyses.**
SV-AUC analysis of 5 μM MLL1 core complex was performed at 25°C with the resulting *c(s)* plot shown in the top two panels. The position of the holo-complex (MWRAD$_2$) at 7.2 *S* is indicated. The range of positions for the unidentified sub-complex peaks X and Y *S* are shown in the dotted rectangles. Since each sub-complex peak represented ~ 5% of the total absorbance, we estimated these sub-complexes to be present in the sample at a concentration of ~0.25 μM. We then mixed stoichiometric amounts of individual subunits for each of the indicated sub-complexes at a final concentration of 0.25 μM and performed SV-AUC at 25°C. The resulting *c(s)* plots for each of the indicated sub-complexes are shown. While unambiguous assignment of the sub-complex peaks could not be determined from direct comparison of *c(s)* plots, the peak positions can serve as powerful restraints in Bayesian analyses of putative assembly schemes. The position of the largest peak in each sub-complex run was used as a prior expectation in the Bayesian analyses reported in Fig. 5. The peak positions of each sub-complex or subunit at 5, 1 and 0.25 μM are summarized in Table S2.

### Figure S2 – Namitz, Tan and Cosgrove

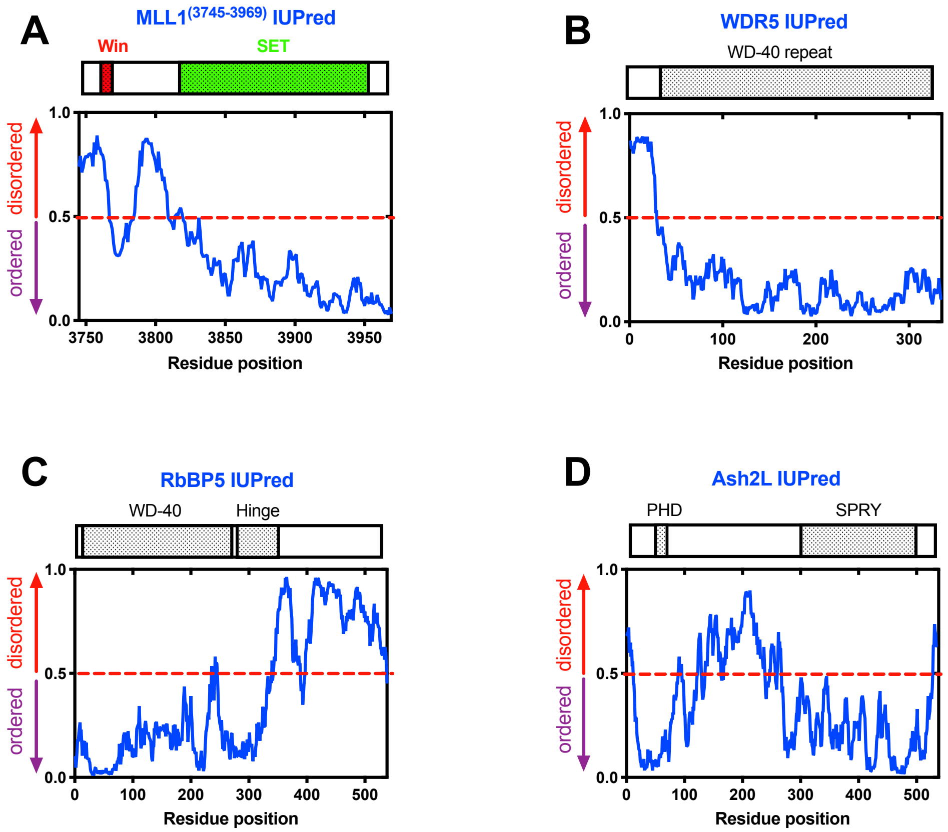


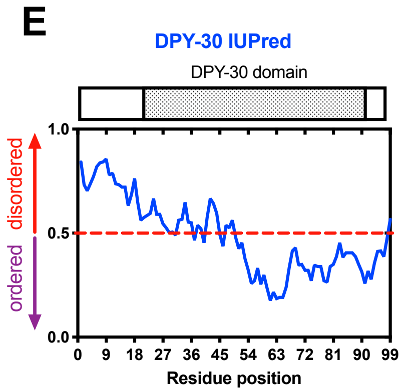


**Figure S2: Each MLL1 core complex subunit has regions of predicted intrinsically disordered amino acid sequences.** IUPred disorder predictions (71) for individual MLL1 core complex subunits: MLL1$^{(3745-3969)}$ (A), WDR5 (B), RbBP5 (C), Ash2L (D), and DPY-30 (E). All $WRAD_2$ components were full-length proteins. Uniprot sub-domain boundaries are shown in the schematic above each panel and are summarized in Table S7.

**Figure S3 – Namitz, Tan and Cosgrove**

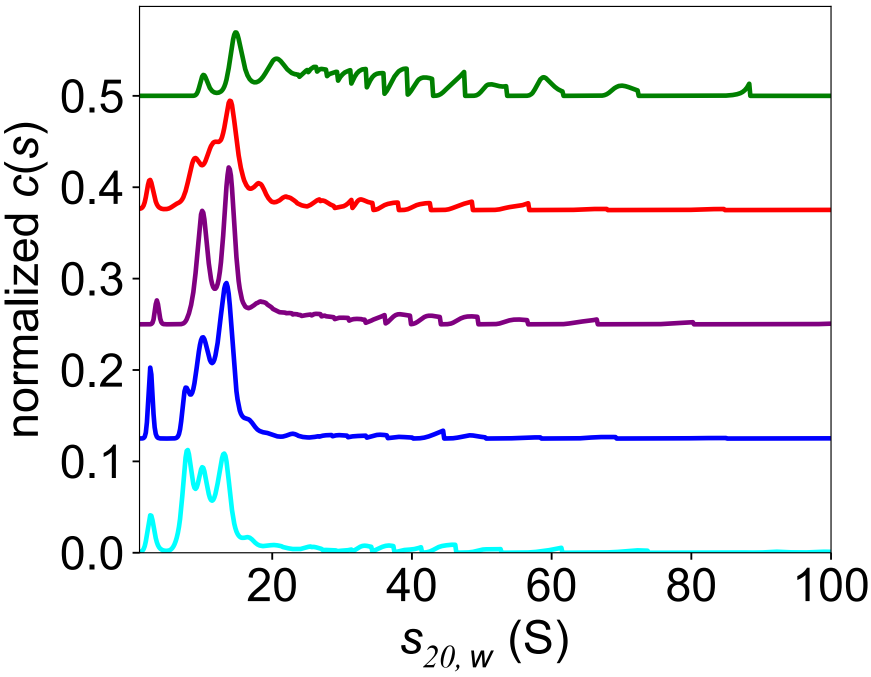


**Figure S3: Concentration dependence of MLL1 core complex oligomerization at low ionic strength.** MLL1 core complex was dialyzed against buffer 3 (with no NaCl), with three changes. SV-AUC runs were conducted with 0.25 (cyan), 0.5 (blue), 0.75 (purple), 1.0 (red) and 5.0 μM (green) $MWRAD_2$ at 25°C. c(s) plots were overlaid, and each was normalized for total integrated area.

**Figure S4 – Namitz, Tan and Cosgrove**

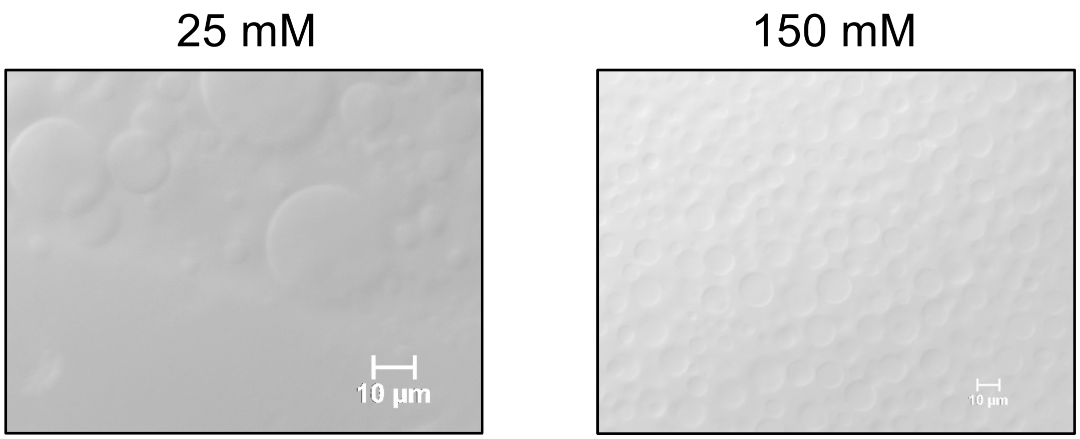


**Figure S4:** $MWRAD_2$ phase separation occurs at both sub-physiological (left panel) and physiological (right panel) [NaCl] in the presence of 7% Dextran.

**Figure S5 – Namitz, Tan and Cosgrove**

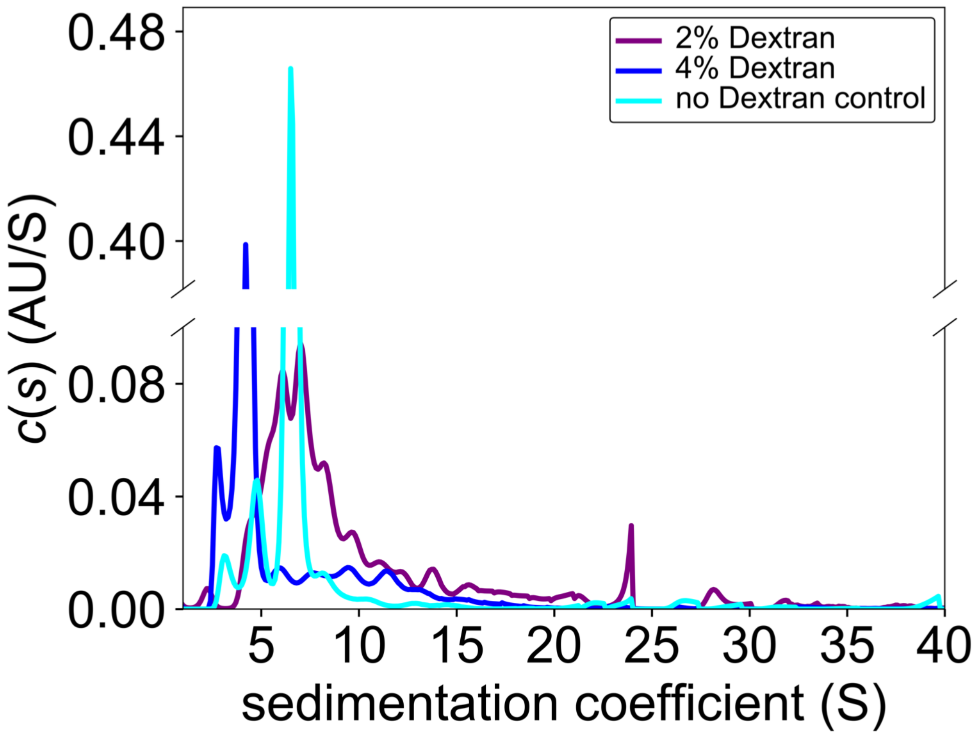


**Figure S5: Dextran induces MLL1 core complex oligomerization at physiological ionic strength that is associated with phase separation.** c(s) distributions of 5 μM $MWRAD_2$ in reaction buffer with 150 mM NaCl at 25°C in absence (cyan), or presence of 2% (purple) and 4% (blue) (w/v) Dextran. Note, the *c(s)* profiles were uncorrected for differences in density and viscosity, which accounts for the slower sedimentation in the 4% Dextran sample, and are shown to illustrate the polydispersity induced by dextran.

**Figure S6 – Namitz, Tan and Cosgrove**

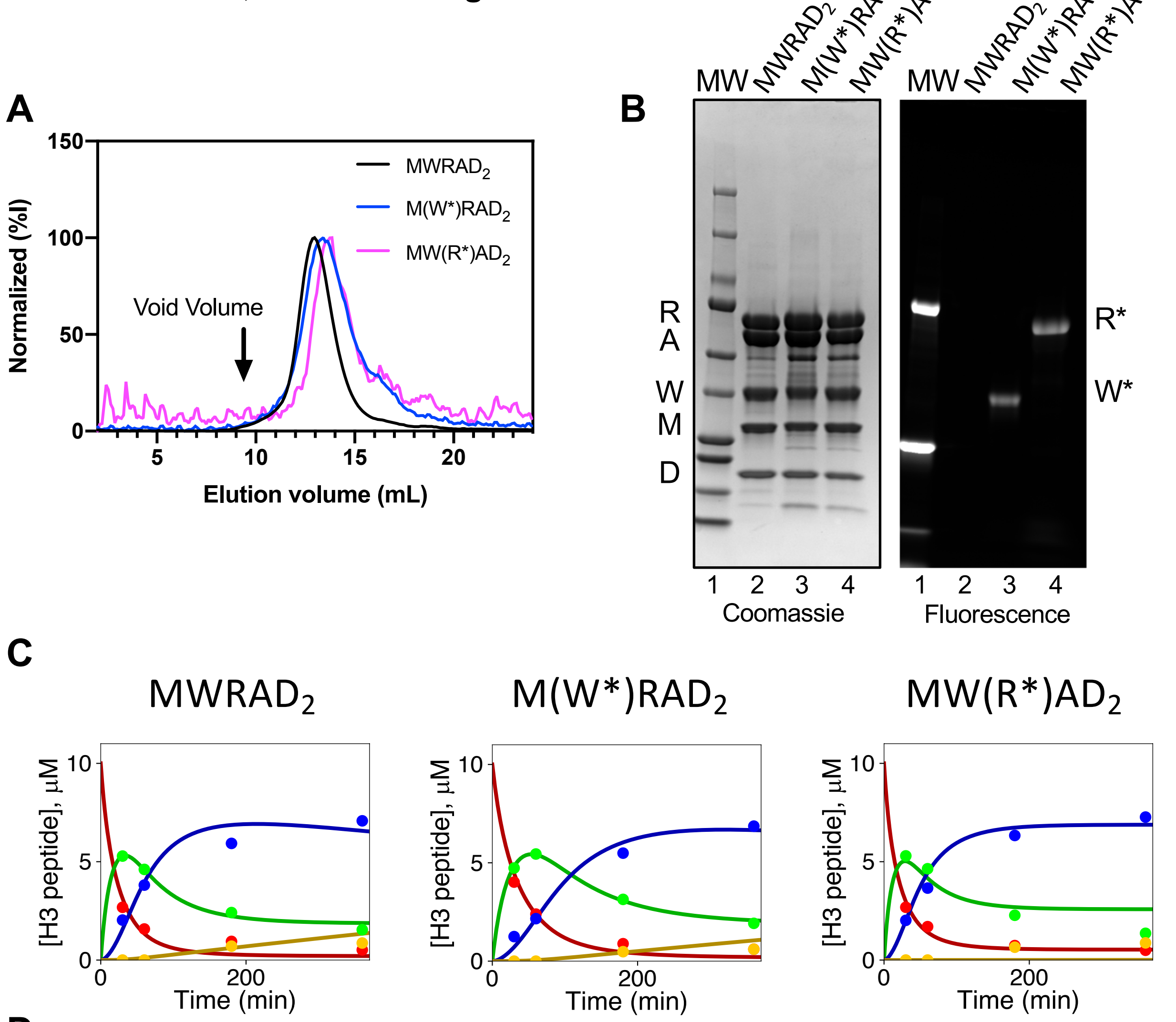


D

| Complex | $k_{me1}$ | $k_{me2}$ | $k_{me3}$ | $k_{inact}$ |
|---|---|---|---|---|
| $MWRAD_2$ | 0.16 (0.02) | 0.12 (0.02) | 0.001 (0.002) | 0.11 (0.03) |
| M(W*)$RAD_2$ | 0.1 (0.01) | 0.07 (0.01) | 0.0009 (0.0008) | 0.07 (0.01) |
| MW(R*)$AD_2$ | 0.19 (0.02) | 0.16 (0.04) | 0.000005 (ND) | 0.17 (0.04) |

**Figure S6: Assembly of fluorescently-labeled MLL1 core complexes.** Representative subunits from the MW or $RAD_2$ sub-complexes were chosen to be N-terminally-labeled with AlexaFluor™ 488. WDR5 (W*) or RbBP5 (R*) were labeled as described in Methods and assembled with the other unlabeled complex subunits and purified by SEC (A). The elution profiles show that the holo-complexes assembled with W* (MW*$RAD_2$) or R* (MWR*$AD_2$) have similar elution profiles to that of the unlabeled complex ($MWRAD_2$). (B) SDS-PAGE of purified complexes visualized by Coomassie blue staining (left panel) or fluorescence imaging (right panel). (C) Reaction progress curves from MALDI-TOF methyltransferase assays comparing the enzymatic activity of unlabeled and labeled complexes at a concentration of 5 μM and 25°C. (D) Summary of pseudo-first order rate constants (S.E.) for unlabeled and labeled complexes.

**Figure S7 – Namitz, Tan and Cosgrove**

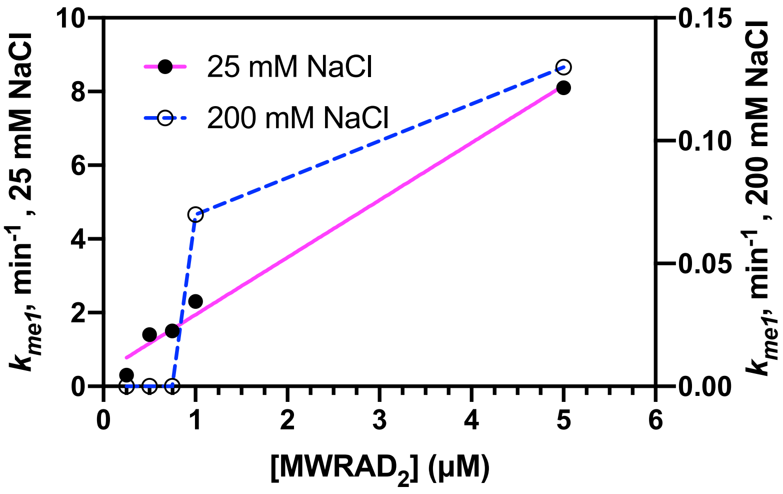


**Figure S7: Comparison of the effect of ionic strength on the concentration dependence of MWRAD$_2$ catalytic activity at 37°C.** Closed circles show $k_{me1}$ rates at low ionic strength (25 mM NaCl), which were plotted on the scale shown on the left. Open circles show $k_{me1}$ rates at high ionic strength (200mM NaCl), which were plotted on the scale on the right. The low ionic strength reaction shows linear dependence over the concentration range (magenta line) (slope = 1.6, $R^2$ = 0.98), whereas the high ionic strength reaction does not (blue line dashed line), primarily due to high rates of irreversible enzyme inactivation at the lower concentrations.

**Figure S8 – Namitz, Tan and Cosgrove**

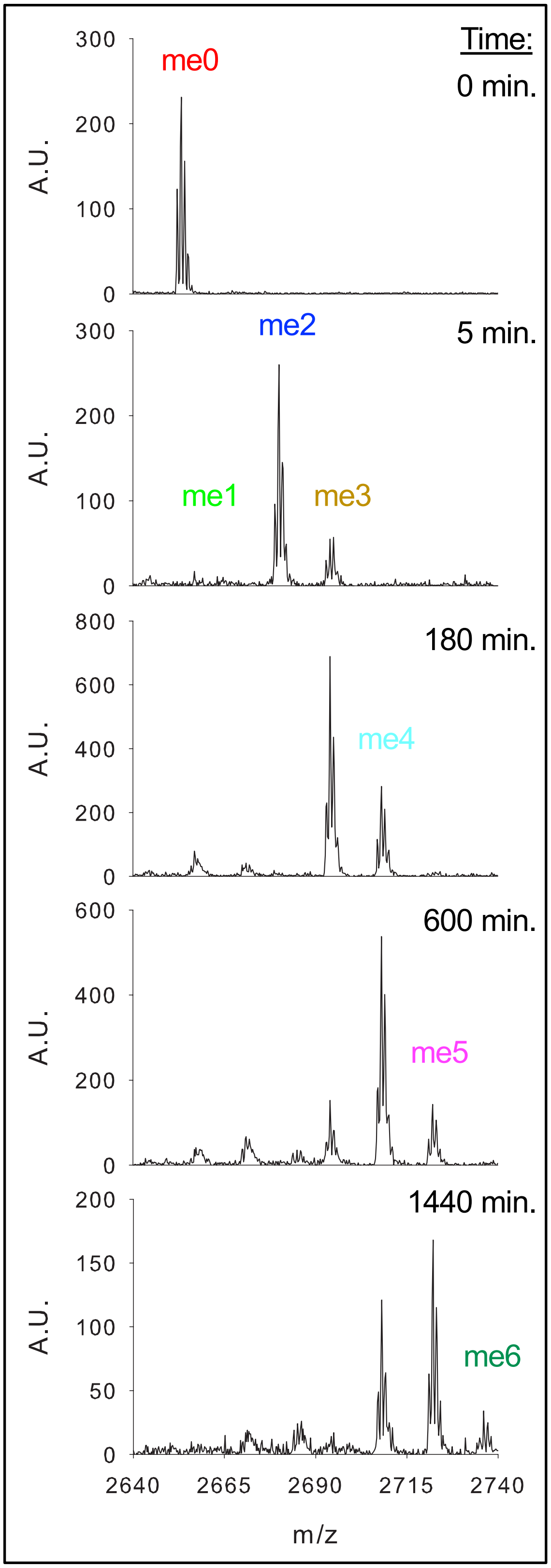


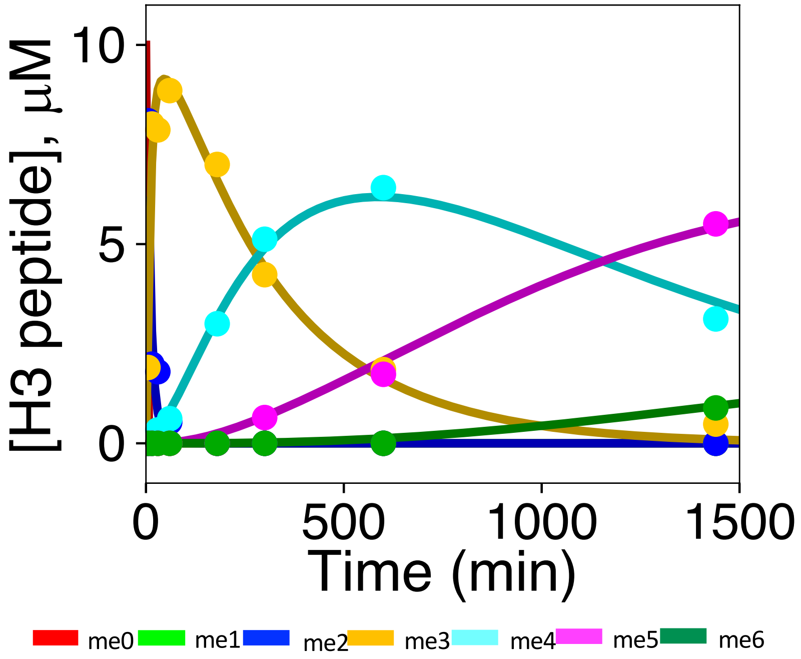


**Figure S8: Up to six methylation events are observed on the same peptide in low ionic strength methyltransferase assays.** Methyltransferase assays were conducted as described in methods in reaction buffer containing 25 mM NaCl. (A) MALDI TOF spectra of time points with each 14 Da shift is indicated. (B) Calculated concentrations of each peptide species were plotted with colors corresponding to that of the labeled peaks in (A).

#### Figure S9 – Namitz, Tan and Cosgrove

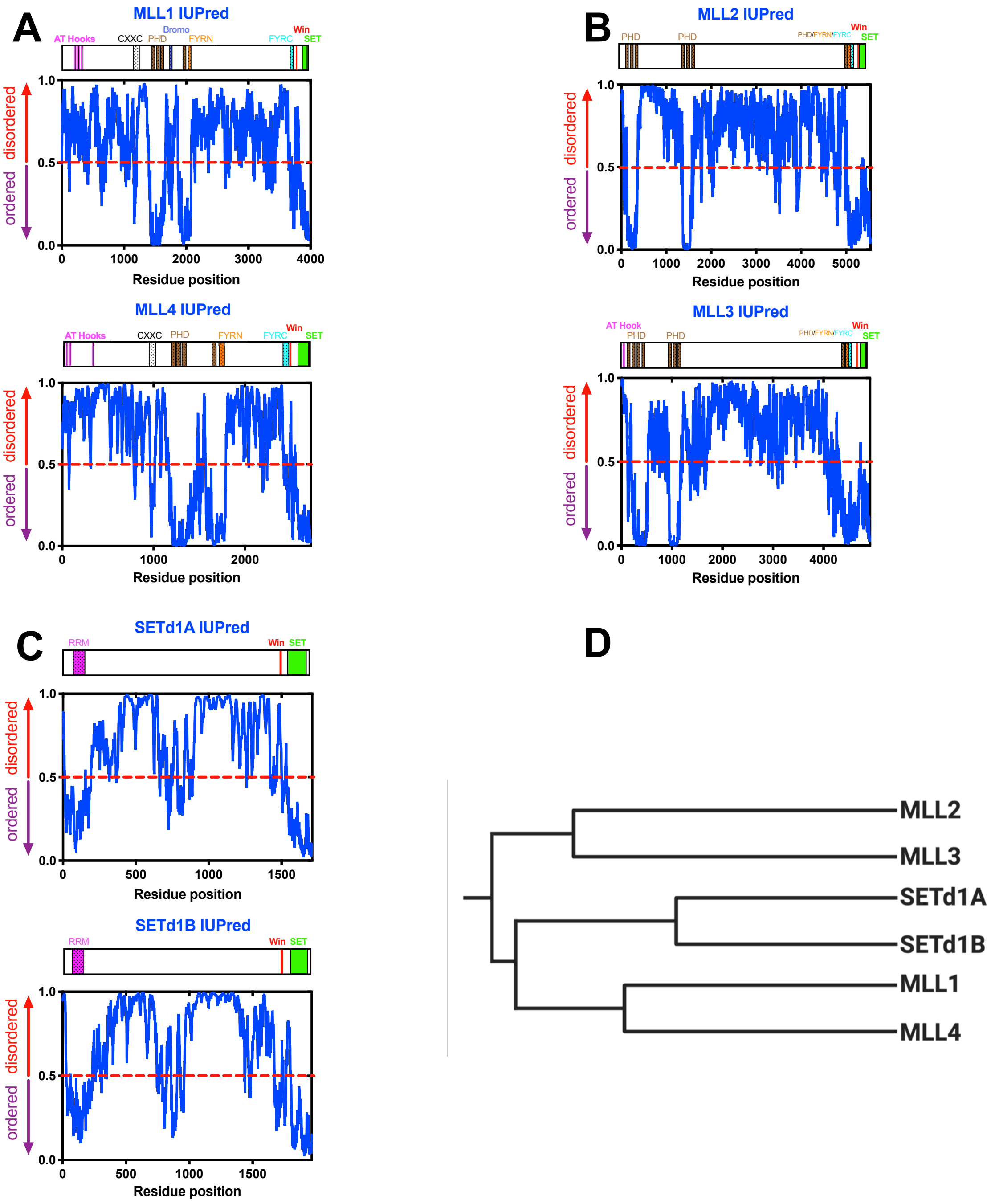


**Figure S9: Each human SET1 family member has large predicted intrinsically disordered regions.** IUPred disorder prediction for (A), full-length Human MLL1 (Uniprot # Q03164) (top) and MLL4 (Uniprot # Q9UMN6) (bottom); (B), MLL2 (Uniprot # O14686) (top) and MLL3 (Uniprot # Q8NEZ4) (bottom); (C), SETd1A (Uniprot # O15047) (top) and SETd1B (Uniprot # Q9UPS6) (bottom). Uniprot sub-domain boundaries are shown in the schematic above each panel and are summarized in Table S6. Note the similarities between the structured domains and linker lengths among the family members from the different phylogenetic clades (MLL1 and MLL4), (MLL2 and MLL3) and (SETd1A and SETd1B). (D) Cladogram showing evolutionary relationships among SET1 family members (created with BioRender.com).

**Movie S1 – Namitz, Tan and Cosgrove**

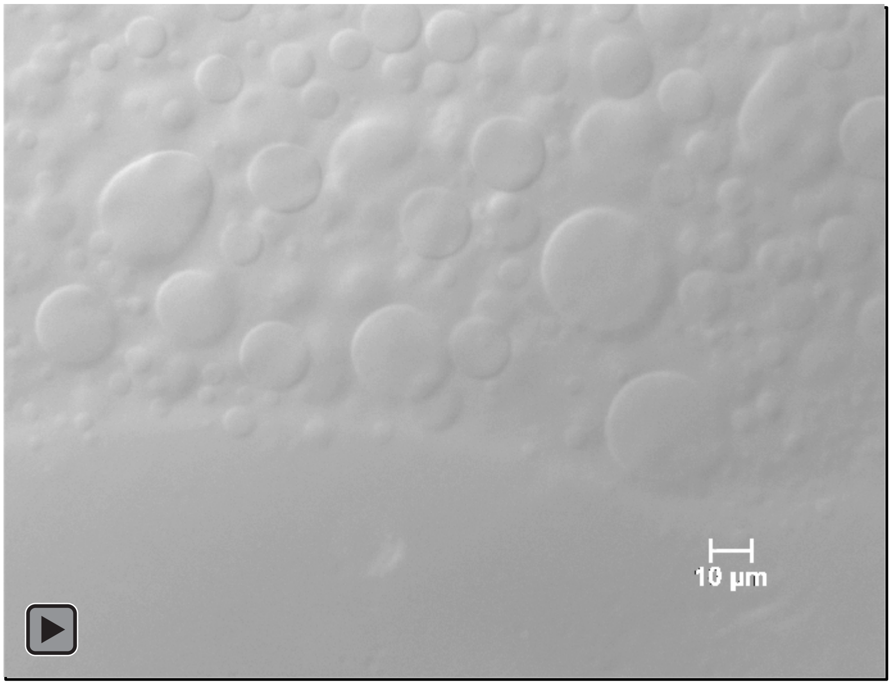


**Movie S1: DIC microscopy movie of 5 µM $MWRAD_2$ in reaction buffer and 7% Dextran.**

7% Dextran – 5µM M(W*)$RAD_2$

**Movie S2: Fluorescence microscopy of M(W*)$RAD_2$ shows localization to phase condensates.** Movie of M(W*)$RAD_2$ from which Figure 9E is taken.

**Movie S3 – Namitz, Tan and Cosgrove**

7% Dextran – 5µM MW(R*)$AD_2$

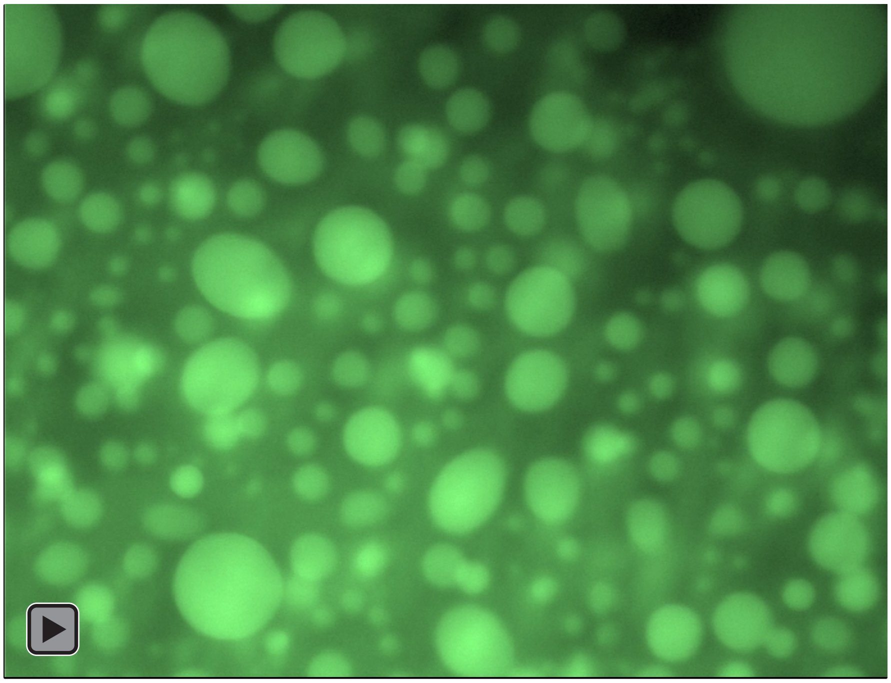

**Movie S3: fluorescence microscopy of MW(R*)$AD_2$ shows localization to phase condensates.**

Movie of MW(R*)$AD_2$ from which Figure 9F is taken.

**Table S1: Summary of the amount of holo-MLL1 core complex from SV-AUC experiments at the indicated concentrations and temperatures.**[a]

| [MWRAD$_2$] (μM) | 5°C | 10°C | 15°C | 20°C | 25°C | 30°C | 37°C |
|---|---|---|---|---|---|---|---|
| 0.25 | 83 (3.0) | 79 (2.0) | 63 (9.0) | 56 (3.0) | 48 (1.0) | 8 (0.7) | 3.0 (2.0) |
| 0.5 | 86 (0.6) | 88 (3.0) | 85 (1.0) | 81 (2.0) | 68 (5.0) | 43 (2.0) | 1.0 (0.3) |
| 0.75 | 91 (0.7) | 90 (0.4) | 89 (0.4) | 86 (3.0) | 78 (0.0) | 55 (1.0) | 0.9 (0.1) |
| 1.0 | 90 (0.5) | 83 (3.0) | 82 (2.0) | 81 (1.0) | 76 (0.5) | 56 (5.0) | 1.3 (0.5) |
| 5.0 | 88 (0.3) | 92 (0.4) | 81 (4.0) | 88 (2.0) | 91 (0.4) | 57 (6.0) | 2.1 (2.0) |

[a] Each value represents the mean percent (±S.D.) of holo-MLL1 core complex signal sedimenting between 6.8-7.6 *S* out of the total integrated signal of all species at each of the indicated loading concentrations and temperatures. Each experimental condition was conducted in duplicate or triplicate.

**Table S2: Summary of *S*-values for MLL1-core complex subunits and sub-complexes from SV-AUC experiments at 25° C.**

| Protein | $s_{20,w}$ (5 μM) | $s_{20,w}$ (1 μM) | $s_{20,w}$ (0.25 μM) |
|---|---|---|---|
| MWRAD$_2$ | 7.2 | 7.2 | 6.9 |
| WRAD$_2$ | 5.5 | 5.2 | 4.4 |
| RAD$_2$ | 5.2 | 5.2 | 4.4 |
| MWR | 4.8 | 3.6 | 3.3 |
| WR | 4.1 | 3.3 | 3.2 |
| AD$_2$ | 4.1 | 4.1 | 4.1 |
| MW | 3.9 | 3.8 | 3.0 |
| M | 2.3 | 2.3 | 2.3 |
| W | 3.2 | 3.2 | 3.2 |
| R | 3.4 | 3.4 | 3.4 |
| A | 3.7 | 3.7 | 3.7 |
| D$_2$ | 1.9 | 1.9 | 1.9 |

**Table S3: Liquid-Liquid Phase Separation (LLPS) Prediction scores for the SET1 H3K4 methyltransferase family**

| Protein | PScore[a] | CatGRANULE[b] |
|---|---|---|
| MLL1 | 4.63 | 1.58 |
| MLL2 | 5.49 | 1.45 |
| MLL3 | 4.64 | 1.37 |
| MLL4 | 5.25 | 1.52 |
| SETd1A | 5.00 | 1.08 |
| SETd1B | 4.25 | 0.95 |

[a]: Per Vernon et al. (106), PScore scores ~4.0 or larger are considered strong candidates for phase separation.
[b]: Per Bolognesi et al. (107), CatGRANULE scores ~1 or larger are enriched in granule-forming (LLPS) proteins.

**Table S4: Liquid-Liquid Phase Separation (LLPS) Prediction scores for the MLL1 core complex individual subunits, sub-complexes and $MWRAD_2$**

| Protein | PScore[a] | CatGRANULE[b] |
|---|---|---|
| MLL1 | 0.54 | 0.32 |
| WDR5 | 0.11 | 0.44 |
| RbBP5 | 0.67 | 0.80 |
| Ash2L | 1.80 | 1.07 |
| MW | 0.58 | 0.76 |
| $RAD_2$ | 1.76 | 1.23 |
| $MWRAD_2$ | 1.78 | 1.36 |

[a]: Per Vernon et al. (106), PScores >1 show possible candidates for LLPS.
[b]: Per Bolognesi et al. (107), CatGRANULE scores ~1 or larger are enriched in granule-forming (LLPS) proteins.

**Table S5: Summary of density, viscosity and partial specific volume estimates.[a]**

| Temperature (°C) | Density | Viscosity | Partial Specific Volume ($v_{bar}$) |
|---|---|---|---|
| 5 | 1.0129 | 0.01569 | 0.724 |
| 10 | 1.0126 | 0.01351 | 0.726 |
| 15 | 1.0120 | 0.01176 | 0.728 |
| 20 | 1.0111 | 0.01035 | 0.730 |
| 25 | 1.0099 | 0.00920 | 0.733 |
| 30 | 1.0085 | 0.00824 | 0.735 |
| 37 | 1.0062 | 0.00714 | 0.738 |

[a] Density and viscosity values were derived from inputting buffer components into SEDNTERP at the indicated temperatures. $v_{bar}$ at each temperature was calculated from the $MWRAD_2$ amino acid sequence using SEDNTERP.

**Table S6: Sub-domain boundaries for human SET1/MLL family histone methyltransferases.[a]**

| (Uniprot #) | MLL1 (Q03164) | MLL2 (O14686) | MLL3 (Q8NEZ4) | MLL4 (Q9UMN6) | SETd1A (O15047) | SETd1B (Q9UPS6) |
|---|---|---|---|---|---|---|
| **Domain[b]** | | | | | | |
| PHD | 1431 – 1482[b]<br>1479 – 1533<br>1566 – 1627<br>1931 – 1978 | 170 – 218<br>226 – 276<br>273 – 323<br>1377 – 1430<br>1427 – 1477<br>1504 – 1559<br>5090 – 5137 | 283 – 331<br>341 – 391<br>388 – 438<br>464 – 520<br>957 – 1010<br>1007 – 1057<br>1084 – 1139<br>4460 – 4507 | 1201 – 1252<br>1249 – 1303<br>1335 – 1396<br>1639 – 1686 | | |
| Bromo | 1703 – 1748 | | | | | |
| AT Hooks | 169 – 180<br>217 – 227<br>301 – 309 | | 34 - 46 | 37 – 44<br>110 – 117<br>357 – 365 | | |
| CXXC | 1147 – 1195 | | | 959 – 1006 | | |
| FYRN | 2018 – 2074 | 5175 – 5235 | 4545 – 4605 | 1727 – 1783 | | |
| FYRC | 3666 – 3747 | 5236 – 5321 | 4606 – 4691 | 2411 – 2492 | | |
| RRM | | | | | 84 – 172 | 93 – 181 |
| Win | 3762 – 3767 | 5337 – 5342 | 4707 – 4712 | 2508 – 2513 | 1492 – 1497 | 1745 – 1750 |
| SET | 3829 – 3945 | 5397 – 5513 | 4895 – 4911 | 2575 – 2691 | 1568 – 1685 | 1827 – 1944 |
| Post-SET | 3953 – 3969 | 5521 – 5537 | 4895 – 4911 | 2699 – 2715 | 1691 – 1707 | 1950 – 1966 |

[a] Protein names and Uniprot # are listed in the topmost row.
[b]Amino acid residue range marking the beginning and end of each sub-domain, which were compiled from Uniprot.

**Table S7: Sub-domain boundaries for $WRAD_2$ sub-complex members.**[a]

| (Uniprot #) | WDR5 (P61964) | RbBP5 (Q15291) | Ash2L (Q9UBL3-3) | DPY30 (Q9C005) |
|---|---|---|---|---|
| **Domain**[b] | | | | |
| WD40 | 43 – 333 | 22 – 331 | | |
| Hinge | | 330 – 366 | | |
| PHD | | | 23 – 56 | |
| SPRY | | | 266 – 489 | |
| DD | | | | 45 – 99 |

[a] Protein names and Uniprot # are listed in the topmost row.

[b] Amino acid residue range marking the beginning and end of each sub-domain, which were compiled from Uniprot.
